# A scenario-guided strategy for the future management of biological invasions

**DOI:** 10.1101/2022.09.07.506838

**Authors:** Núria Roura-Pascual, Wolf-Christian Saul, Cristian Pérez-Granados, Lucas Rutting, Garry D. Peterson, Guillaume Latombe, Franz Essl, Tim Adriaens, David C. Aldridge, Sven Bacher, Rubén Bernardo-Madrid, Lluís Brotons, François Diaz, Belinda Gallardo, Piero Genovesi, Marina Golivets, Pablo González-Moreno, Marcus Hall, Petra Kutlesa, Bernd Lenzner, Chunlong Liu, Konrad Pagitz, Teresa Pastor, Wolfgang Rabitsch, Peter Robertson, Helen E. Roy, Hanno Seebens, Wojciech Solarz, Uwe Starfinger, Rob Tanner, Montserrat Vilà, Brian Leung, Carla Garcia-Lozano, Jonathan M. Jeschke

**Affiliations:** Departament de Ciències Ambientals, Facultat de Ciències, Universitat de Girona, 17003, Girona, Catalonia; Institute of Biology, Freie Universität Berlin, Königin-Luise-Str. 1-3, 14195 Berlin, Germany; Leibniz Institute of Freshwater Ecology and Inland Fisheries (IGB), Müggelseedamm 301, 12587 Berlin, Germany; Berlin-Brandenburg Institute of Advanced Biodiversity Research (BBIB), Königin-Luise-Str. 2-4, 14195 Berlin, Germany; Department of Ecology, University of Alicante, Alicante, 03690, Spain; Copernicus Institute of Sustainable Development, University of Utrecht, Princetonlaan 8a, 3584 CB Utrecht, The Netherlands; Stockholm Resilience Centre, Stockholm University, Kräftriket 2B, 106 91 Stockholm Sweden; BioInvasions, Global Change, Macroecology Group, Department of Botany and Biodiversity Research, University of Vienna, Rennweg 14, 1030 Vienna, Austria; Institute of Evolutionary Biology, The University of Edinburgh, King’s Buildings, Edinburgh EH9 3FL, United Kingdom; Research Institute for Nature and Forest (INBO), Herman Teirlinckgebouw Havenlaan 88 bus 73, B-1000 Brussels, Belgium; Department of Zoology, The David Attenborough Building, University of Cambridge, Downing Street, Cambridge, CB2 3QZ, United Kingdom; BioRISC, St. Catharine’s College, Cambridge, CB2 1RL, United Kingdom; Department of Biology, University of Fribourg, Chemin du Musée 15, 1700 Fribourg, Switzerland; Estación Biológica de Doñana (EBD-CSIC), Av. Américo Vespucio 26, 41092, Sevilla, Spain; CREAF, ES-08193, Bellaterra, Spain; CSIC, ES-08193, Cerdanyola del Vallès; CTFC, ES-25280, Solsona, Spain; Preparedness and Resilience Department, World Organisation for Animal Health, 12 rue de Prony, 75017 Paris, France; Instituto Pirenaico de Ecología, CSIC, Avda. Montañana 1005, 50192 Zaragoza, Spain; Institute for Environmental Protection and Research (ISPRA), Rome, Italy; Chair IUCN/SSC Invasive Species Specialist Group, Rome, Italy; Department of Community Ecology, Helmholtz Centre for Environmental Research – UFZ, Theodor-Lieser-Str. 4, 06120 Halle, Germany; Department of Forest Engineering, DendrodatLab- ERSAF. University of Cordoba. Campus de Rabanales, Crta. IV, km. 396, E-14071 Córdoba, Spain; CABI, Bakeham Lane, Egham, Surrey TW20 9TY, United Kingdom; Institute of Evolutionary Biology and Environmental Studies, University of Zurich, Winterthurerstrasse 190, 8057, Zurich, Switzerland; Institute for Environment and Nature, Ministry of Economy and Sustainable Development, Zagreb, Croatia; Institute of Hydrobiology, Chinese Academy of Sciences, Wuhan, China; Department of Botany, University of Innsbruck, Sternwartestraße 15, 6020 Innsbruck, Austria; EUROPARC Federation, Carretera de l’Església 92, 08017 Barcelona, Spain; Environment Agency Austria, Spittelauer Lände 5, 1090 Wien, Austria; Modelling, Evidence and Policy Group, Newcastle University, NE1 7RU, United Kingdom; UK Centre for Ecology and Hydrology, Benson Lane, Wallingford OX10 8BB, United Kingdom; Senckenberg Biodiversity and Climate Research Centre, Senckenberganlage 25, 60325 Frankfurt, Germany; Institute of Nature Conservation, Polish Academy of Sciences, Al. A. Mickiewicza 33, 31-120 Kraków, Poland; Julius Kühn-Institute, Institute for National and International Plant Health; Messeweg 11-12, 38104 Braunschweig, Germany; European and Mediterranean Plant Protection Organization, 21 boulevard Richard Lenoir, 75011 Paris, France; Department of Plant Biology and Ecology, University of Sevilla, Sevilla, Spain; Department of Biology, McGill University, Montreal, Canada, H3A 1B1

**Keywords:** Biodiversity models, Biosecurity, Environmental and socio-economic impacts, Environmental scenarios, Futures, Global environmental change, Invasive non-native species, Management

## Abstract

Future dynamics of biological invasions are highly uncertain because they depend on multiple environmental, societal and socio-economic drivers. We adopted a qualitative scenario approach to explore the future of invasive alien species (IAS) in Europe and created an overall strategy for their management that considers different plausible future developments. The scenarios and strategy were developed during two online workshops with a multidisciplinary team of experts. First, we downscaled four global scenarios of biological invasions to the European level. Second, we developed a management strategy structured into 19 goals that cover a broad array of IAS-related topics (i.e. policy, research, public awareness and biosecurity), and provided solutions for achieving these goals considering the European scenarios. Third, we identified four interrelated recommendations around which any long-term strategy for managing IAS in Europe can be structured: (i) a European biosecurity regime, (ii) a dedicated communication strategy, (iii) data standardization and management tools, and (iv) a monitoring and assessment system. Finally, we identified the feasibility of the IAS management strategy, finding strong differences among the four scenarios. High levels of technological development, public environmental awareness, and effectiveness of IAS policies facilitated the implementation of the overall management strategy. Together, our results indicate that it is time for a new management of biological invasions in Europe based on a more integrative perspective across sectors and countries to be better prepared for future challenges.

## Introduction

Invasive alien species (IAS) are species that have been introduced by humans beyond their native geographic range and that impact biodiversity and related ecosystem services. IAS are key drivers of global environmental change and strongly contribute to global biodiversity loss. They disrupt ecosystem functioning and affect people through direct health impacts and the damage of agro-ecosystems, cultural landscapes, infrastructure and housing (Potgieter *et al*. 2017; IPBES 2019; Pyšek *et al*. 2020). Further, they produce substantial and increasing economic losses related to direct damages and associated management actions (Vilà *et al*. 2010; Diagne *et al*. 2021).

Despite ongoing efforts in policy, research and management, the number of alien species is still increasing both globally and within Europe with no sign of saturation (Early *et al*. 2016; Seebens *et al*. 2017). In 2014, the European Parliament approved Regulation No. 1143/2014 on the prevention and management of the introduction and spread of IAS, which aims to align and improve the disparate efforts of Member States to counter IAS (Tollington *et al*. 2017). This regulation incorporates recommendations from the Strategic Plan for Biodiversity 2011-2020 of the Convention on Biological Diversity (CBD 2010) to develop early-warning and surveillance systems, action plans to address priority pathways, rapid eradications to prevent establishment, and long-term mitigation and control mechanisms. The implementation of these management recommendations is challenging because of the multiple interacting social-ecological drivers of biological invasions (Dawson *et al*. 2017; Latombe *et al*. 2022), and the uncertainty associated with the future trajectories of societies and global change (Pyšek *et al*. 2020).

Our understanding of IAS patterns and dynamics has improved considerably due to the large-scale mobilisation of information at the global scale (van Kleunen *et al*. 2015; Dyer *et al*. 2017; Biancolini *et al*. 2021). Recent studies have also provided the first quantitative projections of future trajectories of alien species numbers at a continental scale and globally up to 2050 (Sardain *et al*. 2019; Seebens *et al*. 2021). They predict increases in the number of alien species in continental Europe for most taxonomic groups (Seebens *et al*. 2021) and in the risk of marine invasion (Sardain *et al*. 2019). These estimates are based on observed past trends of alien species accumulation or using a reduced set of global economic indicators, neglecting potential shifts in other underlying drivers of alien species movements. They provide a baseline for exploring the future dynamics of biological invasions. However, future number and impacts of IAS are expected to be strongly influenced by the trajectories of multiple environmental, societal and socio-economic drivers over the next decades, which are highly uncertain and therefore difficult to anticipate (Essl *et al*. 2020; Lenzner *et al*. 2020). Quantitative modelling frameworks that incorporate these complex dynamics for predicting biological invasions are still largely absent and would be based on specific assumptions, which would limit the space of plausible and imaginable trajectories. Hence, the development of qualitative scenarios allows for a more open exploration of plausible futures under different environmental and socio-economic trends (Lenzner *et al*. 2019).

Recently, Roura-Pascual *et al*. (2021) developed the first global scenarios for biological invasions. These scenarios are qualitative narratives that explore plausible future trajectories of biological invasions over the next decades and account for uncertainties in social-ecological developments considered critical for IAS on a global scale, giving a stronger focus to biodiversity assets than other available scenarios (e.g. the shared socioeconomic pathways (O’Neill *et al*. 2017)). They consist of 16 scenarios clustered into four contrasting sets of futures, ranging from high to low levels of biological invasions based on technological and trade developments and public environmental awareness Roura-Pascual *et al*. (2021). Qualitative scenarios are instrumental to inform environmental policy making and planning (Wiebe *et al*. 2018), but these global scenarios for biological invasions have never been applied in a continental context or used to inform a specific policy framework.

This study explores qualitative scenarios on the future of biological invasions in Europe and diverse strategies for managing them. The aim is to draft an IAS management strategy to reach an overall vision in the coming decades that is sufficiently robust in the face of critical future uncertainties. This IAS management strategy was developed through a participatory process with a multidisciplinary team consisting of experts in invasion ecology, global change, policy and management, where participants: (i) downscaled four global scenarios of biological invasions (Roura-Pascual *et al*. 2021) to the European level; (ii) developed an overarching management strategy for biological invasions in Europe; (iii) examined the relationship between the different elements of the strategy; and finally (iv) assessed the feasibility of the strategy in the context of the downscaled scenarios and proposed actions to improve it under the challenges posed by each individual scenario. The exact procedure of downscaling the global scenarios to the European level and the relationships between these scenarios is described in another publication. Here, we focus on the development and structure of the IAS management strategy and the assessment of its feasibility under the different scenarios. While our study focuses on Europe, the same procedure can be applied to other geographical regions and at different spatio-temporal scales. Our intention is thus to deliver an overarching guiding framework on how to tackle the long-term management of biological invasions in Europe and elsewhere.

## Methods

### Management strategy development

We adopted a participatory process to develop the management strategy, combining two online workshops (1-2 April and 30 September-2 October 2020) with expert-based internal discussions (Figure 1, WebPanel 1). The structure of the process was based upon previous experiences of the team members with in-person and online scenario workshops (e.g., Roura-Pascual *et al*. 2021). A total of 35 persons participated in the strategy development, with most participants attending both workshops. Attendees represented 12 European countries and three stakeholder groups (i.e. public administration, NGO/interest groups, academia) and were selected with the explicit aim to include a high diversity of professional expertises, perspectives and disciplines. Expertise ranged from hands-on managers and policy-makers (8 participants) to invasion ecologists (23) and from global change and environmental history experts (3) to scenario specialists (1) (WebTable 1). An expert on scenario analysis coordinated the entire process and five additional participants facilitated the breakout groups. Together, they formed the team of workshop facilitators. Other members were assigned to breakout groups and provided their expertise in different workshop sessions.

**Figure 1.**
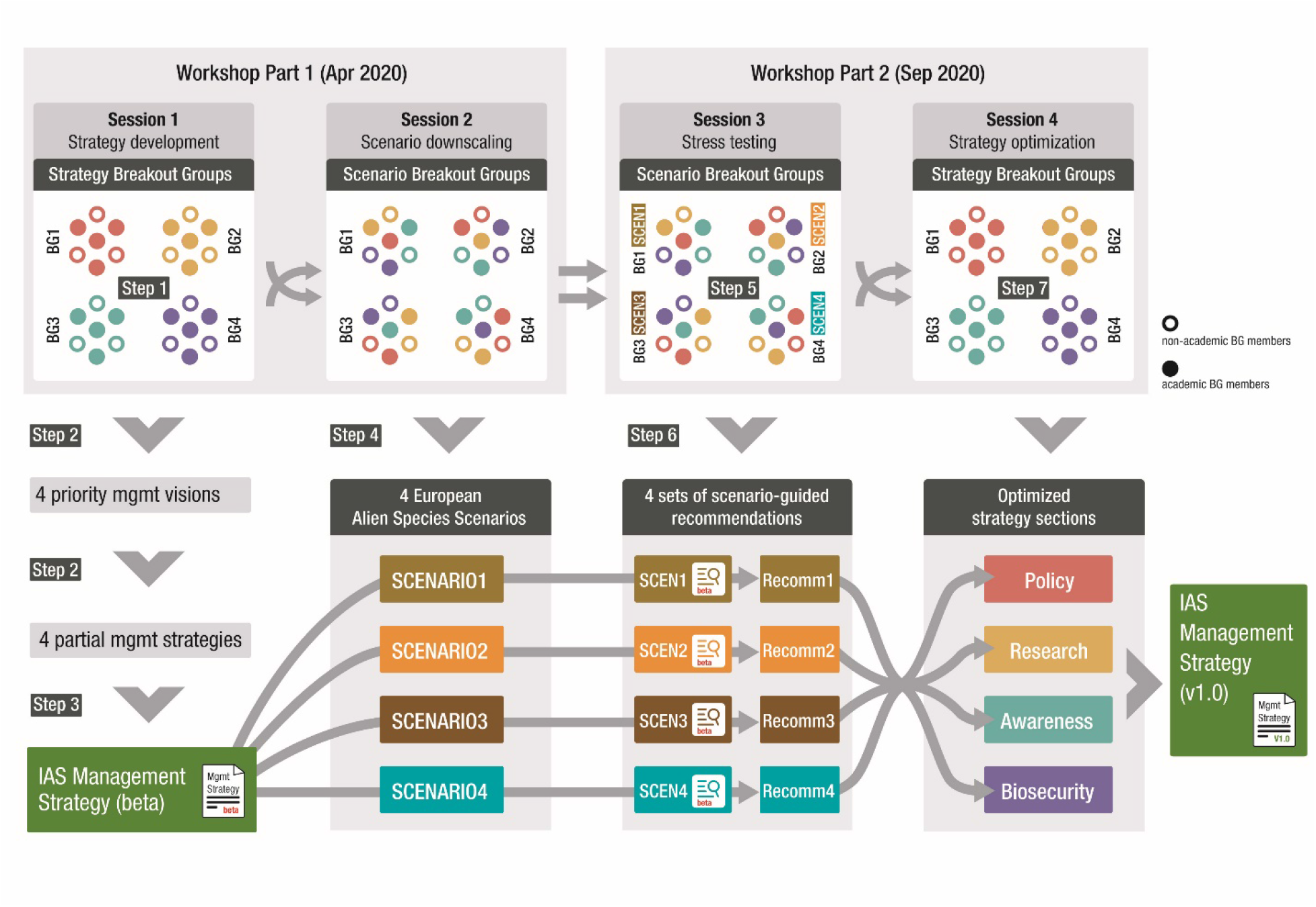
Workflow diagram of the participatory process used to explore the future management of biological invasions in Europe, which resulted in an IAS management strategy and four qualitative scenarios describing potential future developments (abbreviated SCEN). For a more detailed description of the steps, see Methods and WebPanel 1.

The participatory process consisted of seven steps. The first four steps were developed during the first online workshop, while the last three during the second one (Figure 1, WebPanel 1). In Step 1, the participants were divided into four *strategy* breakout groups to formulate general visions (i.e. general objectives) for IAS management in the coming decades in Europe. In Step 2, the visions were collected and presented to the entire group of participants, who then voted for the four visions considered to be of highest importance. In Step 3, each of the four most voted visions was assigned to a strategy breakout group, each of which developed a preliminary management strategy aimed to reach its assigned vision. The four resulting partial management strategies were subsequently combined into one overall management vision and strategy (‘beta version’) composed of various goals and actions. The goals refer to specific management aspects identified as important by the workshop participants and can be conveniently grouped into four categories: Policy, Research, Public awareness, and Biosecurity. Actions describe the steps required to reach each goal.

In Step 4, participants were reshuffled into *scenario* breakout groups (each comprising members of all previous strategy breakout groups). These groups reframed and downscaled four previously developed global scenarios for biological invasions (Roura-Pascual *et al*. 2021) to the European level, considering contextual conditions with regard to social, political, economic and environmental developments in Europe. Out of sixteen global scenarios available, four were selected to be downscaled to the European level. The four scenarios were selected as being representative of the four main clusters of scenarios identified in Roura-Pascual *et al*. (2021) and thus cover the full range of global scenarios describing differing future trajectories for IAS with specific challenges and opportunities.

During the second workshop, Step 5 started with revisiting and fine-tuning the four newly developed European scenarios. Afterwards, each scenario breakout group identified weaknesses and strengths of the overall management strategy (‘stress test’) by assessing which actions (and consequently goals) of the strategy would likely be successful or not in the context of each scenario, i.e. ranked the strategy’s feasibility into feasible, partially feasible, or not feasible. An action assessed as not feasible or partially feasible was considered an action to be improved to make it feasible. Based on this, in Step 6 the groups formulated recommendations for improving these actions and therefore the overall strategy. Finally, in Step 7, participants reconvened into strategy breakout groups and each group was responsible for a different category of goals (Policy, Research, Public awareness, Biosecurity). Each group discussed the proposed recommendations and amended the objectives/actions accordingly, which were then integrated into one final overall management strategy (*v1*.*0*; Step 7, Figure 1).

### Management strategy analysis

We identified associations between the different goals of the IAS management strategy using a two-step process. First, the workshop facilitators captured the essence of each goal by using keywords. Each facilitator was in charge of a specific category of goals (Policy, Research, Public awareness, Biosecurity) and was responsible for: (i) characterizing the goals of the selected category, and then (ii) reviewing all goals of the management strategy to identify associations between the goals of the focal category with goals of the other three categories. An association is loosely defined here as any justifiable direct relationship between two goals of the strategy, without necessarily implying a causal relationship. Second, the facilitators from the first step focused solely on their own category and revised the associations proposed by the other facilitators on the focal category. In case of disagreement, a consensual decision was adopted after discussion among the facilitators. At the end of this two-step process, the strategy and the associations between goals were sent to all workshop participants to identify and remediate potential inconsistencies. We used a chord diagram to represent the number of associations between categories of goals and a table to show the associations between goals.

Additionally, we examined the feasibility of the IAS management strategy under the different scenarios. In an online exercise after the workshop, each scenario breakout group revisited the feasibility assessment conducted in Step 6 and repeated the process for the actions (i.e. steps required to reach each goal) of the final management strategy. We converted these feasibility assessments given to each action of the strategy into a numerical value: feasible action = 1; partially feasible action = 0.5; unfeasible action = 0. We averaged these values at the goal level and calculated the mean of the goals’ averaged values at the category level (Policy, Research, Public awareness, and Biosecurity). We plotted the mean (and standard error) feasibility of the different categories under the different scenarios to graphically reveal differences among future developments. Similarly, we plotted the feasibility of the different goals grouped by categories to graphically identify the divergences between scenarios.

## Results and Discussion

### IAS management strategy

Participants to the workshops explored different visions and strategies for the future management of biological invasion in Europe, but they finally agreed to combine them into one overall management vision and strategy because of their similarities. The resulting vision was: “By 2050, the harmful impacts of IAS in Europe (EU-Member States and non-EU states) are substantially reduced compared to today”. This vision is in concordance with ongoing international negotiations on the global post-2020 biodiversity framework of the CBD, which includes a target directed to reduce the introduction and establishment of IAS and their impacts (CBD 2021). Impacts are changes caused by IAS to their ecological and/or socio-economic environment. They integrate the range, abundance and per-capita effects of IAS (Parker *et al*. 1999) and are related to the introduction, establishment and spread of IAS (Robertson *et al*. 2020). They therefore act as surrogates of a wider range of factors. In addition, they are ultimately the reason for targeted IAS management.

The IAS management strategy is structured around 19 goals and a series of actions per goal that are considered relevant for ensuring the achievement of the general vision (WebPanel 2). These goals cover a broad array of IAS issues beyond those strictly related to management and can be grouped into four broad categories: Policy, Research, Public awareness and Biosecurity. Below, we provide a brief presentation of the goals included in each category and indicate the most prominent cross-cutting aspects that emerged within each category. The complete IAS management strategy can be consulted in WebPanel2.

- Policy (goals P1-P6) refers to the competencies of political actors to harmonize and strengthen IAS regulations at the European and country levels (abbreviated P1), the identification and securing of long-term funding for IAS management (P2), horizon scanning of threats and opportunities for management and incorporation into IAS policies (P3), and prioritization at the species (P4), sites (P5) and pathway levels (P6). Each goal focuses on a particular policy aspect, but all of them highlight the need to integrate countries’ policies, priorities and funding at the European level. IAS move across political borders as long as there is suitable habitat available, so managing IAS efficiently requires aligning policy efforts and resources across countries (Keller *et al*. 2011).
- Research (goals R1-R4) includes all aspects concerning scientific expertise, from the creation of research networks (between researchers, but also among stakeholders) (R1) to targeted research on data gaps (R2), critical tools (R3) and other knowledge gaps (R4). These goals together emphasize the importance of conducting collaborative research, not only limited to scientists from different disciplines but also involving stakeholders from different sectors, to establish a standardized terminology/methodologies and open access data/tools/information across Europe. Scientific literature dedicated to applied studies and field experiments that provide management solutions is proportionally low in comparison to theoretical studies (Bayliss *et al*. 2013; Muñoz-Mas *et al*. 2021). The adoption of such a collaborative approach would facilitate responses to management needs and the availability and accessibility of IAS data, which ultimately support a cost-efficient IAS policy and management (Gatto *et al*. 2013).
- Public awareness (goals A1-A3) relates to the development of a communication strategy and platform (A1), targeted funding for raising public awareness (A2) and the active engagement of the general public and stakeholders (A3). Public awareness on IAS management and citizen science programmes are growing, but there are still conflicting ethical views in the general public (e.g. regarding the killing of IAS animals) (Novoa *et al*. 2017) and among managers (e.g. removal of established IAS) (Blaalid *et al*. 2021) or scientists (Shackleton *et al*. 2022). Additionally, the communication between European countries is still inefficient. Public engagement and public awareness should be promoted by a dedicated cross-sectoral communication campaign/platform that facilitates knowledge transfer across Europe. The use of citizen science platforms together with several other strategies for raising public awareness and engaging the public with the IAS problem have been validated as an effective tool to improve IAS management at the country level (Marchante and Marchante 2016; Probert *et al*. 2022).
- Biosecurity (goals B1-B6) encompasses the establishment of a common biosecurity at the European level (B1) and increased international cooperation among institutions and stakeholders (B2), as well as aspects related to monitoring (B3), management (prevention, eradication, control) of IAS (B4 & B5) and subsequent ecological restoration (B6). Decision making applied to IAS is complex and management practices are affected by multiple elements (Dana *et al*. 2019), but there is an impressive body of knowledge and technical information on IAS management accumulated through years of management practices (Scalera *et al*. 2017). Here (as it happens with the policy goals) participants stressed the relevance of fostering collaborations between different stakeholders and coordinating management efforts within Europe and beyond European borders. The coordination of actions requires a continental and international approach (Hulme 2021).

Since this strategy has been developed together with stakeholders across Europe, it provides a general picture of what elements of IAS management are considered important within Europe. Several of these elements (herein considered goals) have already been identified as relevant for managing biological invasions (Piria *et al*. 2017) and are even included in a IAS management framework to standardize management terminology (Robertson *et al*. 2020). Here we organize them in an overall framework to guide action on IAS taking into account future uncertainties and thereby contributing to devising the long-term management of IAS in Europe. Due to the inclusion of elements beyond those that are strictly related to management, this framework also represents a further development of the IAS management framework proposed by Robertson *et al*. (2020).

### Strategy’s associations

Besides the importance of considering multiple and diverse goals in IAS management, some goals and categories of goals are more closely associated with each other (Figure 2). At the category level, most associations are between Biosecurity and Policy (35 of 36 possible connections; 97%), and between Biosecurity and Research (21/24; 87%) (Figure 2a). While the least number of associations exist between Public Awareness and Research (8/12, 67%) and Public Awareness and Biosecurity (12/18, 67%), all categories are connected with each other (Figure 2a). The high connectivity among the different categories highlight the integrative nature of the IAS management strategy and the mutual dependency of its components to ensure the accomplishment of its vision.

**Figure 2.**
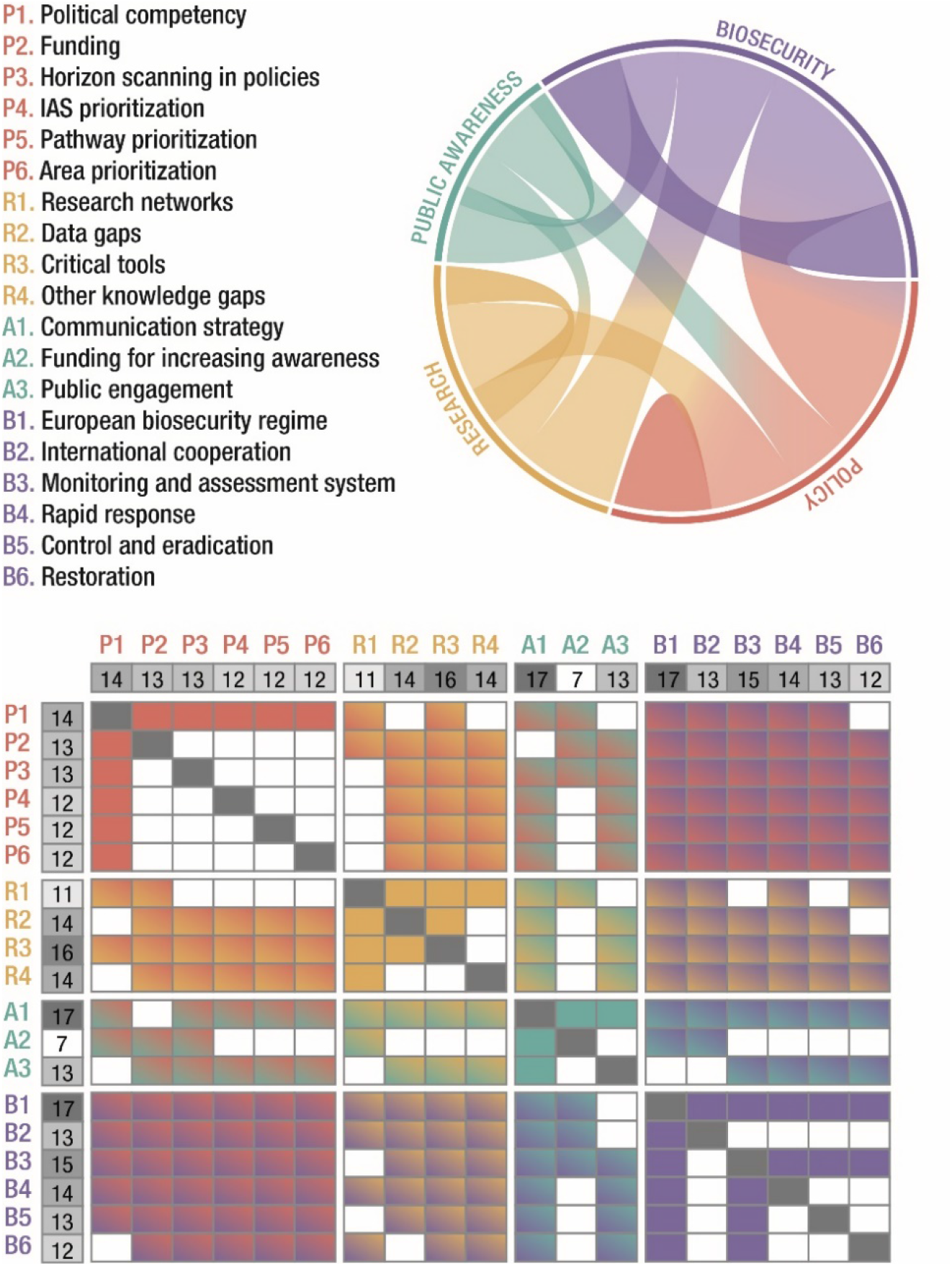
Visualization of associations between categories of goals (chord diagram) and between individual goals (table) of the IAS management strategy. The outer circle of the chord diagram is segmented into the strategy’s four categories: Policy (abbreviated P and coloured in red), Research (R, orange), Public awareness (A, green), and Biosecurity (B, blue). The portion that each segment occupies of the full circle corresponds to the number of associations the respective category has in proportion to the total number of associations identified throughout the entire strategy. The width of the coloured arches connecting two different parts of the circle indicate the number of associations that exist between goals of the two connected categories (or between goals of the same category). In the table, the row and column in shades of grey indicate the total number of goals to which each goal is associated. The central matrix shows the associations between the different goals, with coloured cells indicating an association found between two goals. Cell colours reflect the categories of the goals involved in the association.

At the goal level and irrespectively of the category they belong to, on average each goal is associated to 13 of the other 18 goals (72%). The goals with most associations are: European biosecurity regime (B1) and Communication strategy (A1), followed by Critical tools (R3) and Monitoring and assessment system (B3) (Figure 2b). The least associated goals of the management strategy are Research networks (R1) and Funding for increasing public awareness (A2) (Figure 2b). Although Policy competency (P1) does not appear as one of the most associated goals, it is well-connected to all other Policy goals. These most connected goals represent key elements for the implementation of the overall management strategy and, therefore, deserve particular attention. In Box 1, we present these goals and aforementioned cross-cutting aspects within each category (presented in previous section) as core aspects for cross-cutting recommendations emerging from the overall strategy.

### Strategy’s feasibility

The use of future scenarios for biological invasions improved the feasibility of the goals, and the management strategy as a whole, for any of those futures. Actions required to reach each goal incorporate the recommendations that resulted from the stress-testing process (as described in Step 6). Besides this increased feasibility, our analysis shows that some scenarios (or futures) are more challenging than others (Figure 3). Future scenarios with high levels of technological development, public environmental awareness, and effectiveness of IAS policies that encourage research and biosecurity regarding IAS (like “Technological (Pseudo-)Panacea”) are favourable for the implementation of the overall management strategy across all goals (WebTable 2). More disruptive futures such as those conceiving an isolationist Europe (“Lost in Europe”) will result in considerable difficulties for the strategy implementation. In particular, goals requiring a certain level of coordination across Europe (i.e. all policy-related goals and other key goals, such as the establishment of a European biosecurity regime (B1) and a Communication strategy (A1)) will be extremely difficult to progress under this scenario (WebTable 2).

**Figure 3.**
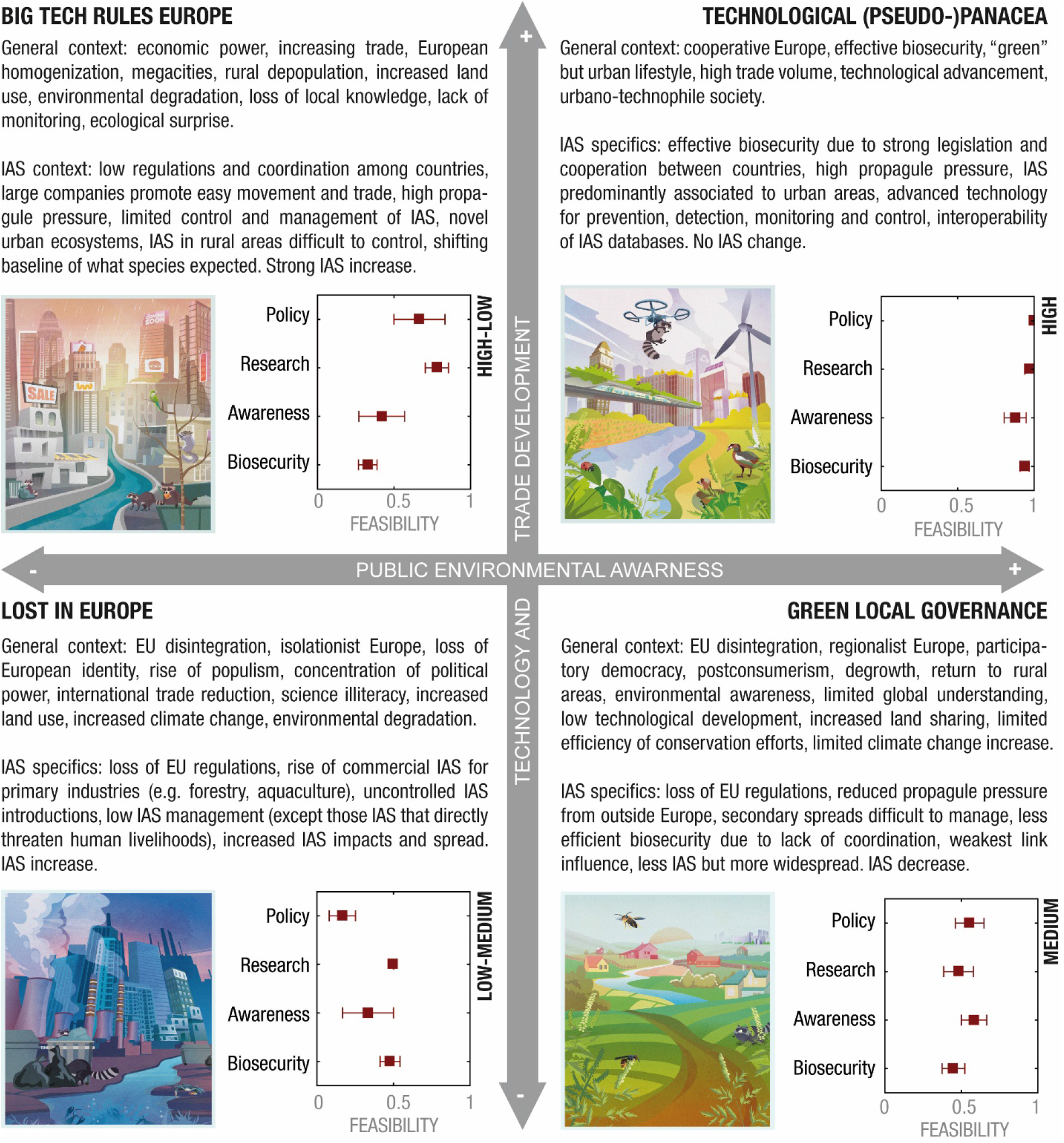
Summaries of the future scenarios for biological invasions in Europe until 2050 and feasibility of the management strategy in each scenario. The four scenarios were downscaled from global scenarios created in an earlier contribution (Roura-Pascual *et al*. 2021). Means and standard errors indicate the average feasibility of the management goals included in the strategy (1 = Feasible; 0.5 = Partially feasible; 0 = Not feasible), as judged by the workshop participants and grouped by categories: Policy, Research, Public awareness and Biosecurity. See WebTable 2 for the feasibility of each individual goal under the different scenario assumptions. IAS refers to invasive alien species.

In between these two extremes are scenarios with a more variable feasibility of the management actions. The different goals (and categories of goals) are affected differently under the different scenarios (Figure 3). Some futures may include a high prominence of economic power and technology that stimulate research and policy development on IAS (“Big Tech Rules Europe”), while other futures may put a strong emphasis on raised public awareness that compensates for technological deficiencies and thus results in a similar feasibility of biosecurity measures (“Green Local Governance”). Irrespective of the scenario, goals related to biosecurity present medium feasibility and therefore suggest that our vision would only be partially achievable. Overall, our results indicate that reducing the impacts of IAS in Europe requires the establishment of an efficient biosecurity programme, but also adequate policy and research contexts that support the development and implementation of the programme and an appropriate communication campaign to facilitate the uptake of the programme by all stakeholders.

## Conclusions

Biological invasions are contingent upon a wide range of environmental, socio-economic and political developments, so any long-term strategy aimed to reduce their impact needs to consider such developments. The IAS management strategy developed herein has been built around 19 policy, research, public awareness and biosecurity goals. The high interdependence found between these goals and categories of goals indicate that successful management of IAS relies on pursuing multiple and diverse goals, particularly those goals that are strongly connected to others. Based on these most connected goals and a series of cross-cutting aspects that emerged within the management strategy, our results highlight that any strategy aimed to prevent and mitigate the impacts of IAS across Europe requires the implementation of four interrelated recommendations: (i) cooperation between countries and stakeholders at the European level, (ii) communication and outreach across sectors, (iii) data standardisation and management tools, and (iv) efficient IAS monitoring and assessment with corresponding management priorities. None of these recommendations will be sufficient on its own, but they have been identified as key elements around which to structure a long-term strategy for managing biological invasions at the European level.

The future feasibility of the IAS management strategy and its respective goals varies across scenarios. Some scenarios are more favorable than others, being scenarios other than the optimal (i.e. with a high level of technological development, public environmental awareness, and effectiveness of IAS policies) less likely to ensure the achievement of the 2050 vision of substantially reducing the impacts of IAS. The scenario process allowed us to examine the management strategy under the lenses of different possible futures and to propose solutions to improve its feasibility. Even though these solutions are not sufficient to achieve the strategy’s vision, their inclusion is an essential step to deliver a long-term strategy that is better prepared to future developments. It is time for a shift in the management of biological invasions in Europe based on a more integrative perspective across sectors and countries.

## Supporting information

Supplementary Matrial (WebTable 1, WebPanel 1, WebTable 2)

Supplementary Material (WebPanel 2)

## Acknowledgements

This research was funded through the 2017-2018 Belmont Forum and BiodivERsA joint call for research proposals, under the BiodivScen ERA-Net COFUND programme, and with the funding organisations AEI, BMBF and FWF [grant numbers AEI PCI2018-092966 (NRP, CPG, CGL) / AEI PCI2018-092939 (MV) / AEI PCI2018-092986 (BG) / BMBF projects 16LC1803A & 16LC1807B (JMJ, WCS) / BMBF 01LC1807A (HS) / FWF project 4011-B32 (FE, GL, BL)]; Swiss National Science Foundation [grant number 31BD30_184114 (SB)]. CPG acknowledges the support from Ministerio de Educación y Formación Profesional through the Beatriz Galindo Fellowship (Beatriz Galindo – Convocatoria 2020). We highly appreciate the participation to the workshops of Spyridon Flevaris, Ingolf Kühn and Jörg Priess, the scenario illustrations of Kris Tsenova, and the constructive comments of anonymous reviewers and the handling editor.

## Box 1

Recommendations for managing biological invasions in European, derived from the most interconnected goals (i.e. B1, A1, R3 and B3) and most relevant cross-cutting aspects emerging from the management strategy. They represent board fundamental principles that lie at the core of the strategy and should lead its implementation.

### Recommendation 1 – European cooperation for a common and effective biosecurity regime

This recommendation (based on goal B1) proposes the strengthening of European cooperation. It advises for the establishment of a dedicated European agency or an intergovernmental agreement furnished with a mandate and resources to regulate, oversee and improve all activities related to IAS management in Europe. This requires a high degree of cooperation and coordination between states and stakeholders. For promoting cooperation between states, European and global international organizations (such as CABI (Centre for Agriculture and Bioscience International; www.cabi.org), EPPO (European and Mediterranean Plant Protection Organization; www.eppo.int), Bern Convention (https://www.coe.int/en/web/bern-convention), CBD (Convention on Biological Diversity; www.cbd.int), IMO (International Maritime Organization; www.imo.org), among others) are expected to provide guidance with respect to IAS policy, prioritization, best practice and management harmonization between countries. They should also support the coordination between EU Member States and non-EU states. To actively integrate different stakeholder perspectives, this agency or agreement should also foster interactions and synergies across sectors and stakeholders, consider regional particularities (e.g. regarding differences in management priorities between European regions or countries), and integrate local knowledge and cultures, including public involvement in decision-making processes. Shared governance and participatory decision-making provide strengthened legitimacy to actions based on such consensus agreements.

### Recommendation 2 – Cross-sectoral IAS communication and outreach strategy

This recommendation (based on goal A1) advises the establishment of a cross-sectoral IAS communication strategy (including a dedicated education curriculum) and a centralized, multilingual communication platform at the European level. Across all sectors and geographic regions in Europe, the communication strategy must help to increase awareness regarding causes and consequences of IAS and their management, while the platform will facilitate knowledge transfer and collaboration. Goals in all categories of the management strategy benefit from principles of good and transparent communication leading to an increased awareness and understanding of IAS-related problems among stakeholders and the general public. Ultimately, this is a prerequisite for strong support and success of management measures.

### Recommendation 3 – Data standardization and management tools

This recommendation (based on R3) consists of regularly addressing critical gaps in tools for IAS impact/risk assessment and their management. Suggested actions include creating and/or improving standard protocols for assessing impacts, pathways and vulnerability of priority areas, conceiving adaptive approaches to guide management decisions, developing novel management techniques, and ultimately ensuring the adoption of these tools at the country and European level, and if applicable even globally. Several other goals (like R2 and R4, among others) call upon the establishment of a European centralized IAS open data portal that facilitates the efficient recording, storing, standardization, continuous updating, peer-reviewing, and broad accessibility of all available information related to IAS and their management in Europe, and allows automated approaches to managing and analysing these big datasets. The European Alien Species Information Network (EASIN; https://alien.jrc.ec.europa.eu/easin) may provide a useful basis for this platform. Standardization and data aggregation on open data platforms (e.g. Global Biodiversity Information Facility) facilitates comparisons (e.g. between species, invasions cases or countries), allows combining data from different sources for greater power of inference, and supports transparent prioritization of management actions.

### Recommendation 4 – IAS monitoring, assessment and management priorities

This recommendation (linked to goal B3) implies the establishment of a comprehensive regime for monitoring and assessing IAS at the European and country levels. Having a sound and comprehensive knowledge of the past, current and future circumstances of the introduction, establishment, and spread of IAS as well as of their (recorded and potential) impacts and past management attempts is a crucial prerequisite for any IAS management strategy to be effective. This information is used or complemented by other goals and actions of the strategy. Particularly, the establishment and implementation of management priorities at the species, site and pathway level is a recurrent element in the management strategy and requires such information. Policy regulations (e.g. the list of invasive alien species of Union concern in Regulation 1143/2014) are an important tool for defining legally binding priorities (see goals P4-P6). On other levels in management practice, however, priorities may change on a more flexible basis, e.g. when managers obtain new data concerning the areas, species or pathways they are managing (see e.g. B4-B6). The latter approach is common in IAS management in practice, but is not necessarily subject to direct political regulatory processes.

