## Supplementary Matrial (WebTable 1, WebPanel 1, WebTable 2) for "A scenario-guided strategy for the future management of biological invasions"

**SUPPORTING INFORMATION**

**WebTable 1.** Participants involved in the development of the management strategy for invasive alien species in Europe, indicating their country and expertise. (*) indicates members of the workshop facilitation team.

| Name | Country | Stakeholder type | Stakeholder expertise |
|---|---|---|---|
| Franz Essl | Austria | Academia | Invasion ecology |
| Bernd Lenzner | Austria | Academia | Invasion ecology |
| Konrad Pagitz | Austria | NGO/Interest group | Policy and management |
| Wolfgang Rabitsch | Austria | Public administration | Invasion ecology |
| Tim Adriaens | Belgium | Academia | Invasion ecology |
| Spyridon Flevaris | Belgium | Public administration | Policy and management |
| Petra Kutlesa | Croatia | Public administration | Policy and management |
| Chunlong Liu | France | Academia | Invasion ecology |
| François Diaz | France | Public administration | Policy and management |
| Rob Tanner | France | Public administration | Policy and management |
| Marina Golivets | Germany | Academia | Invasion ecology |
| Jonathan M. Jeschke* | Germany | Academia | Invasion ecology |
| Ingolf Kühn | Germany | Academia | Invasion ecology |
| Jörg Priess | Germany | Academia | Invasion ecology |
| Wolf-Christian Saul* | Germany | Academia | Invasion ecology |
| Hanno Seebens | Germany | Academia | Invasion ecology |
| Uwe Starfinger | Germany | Academia | Invasion ecology |
| Piero Genovesi | Italy | NGO/Interest group | Policy and management |
| Lucas Rutting* | Netherlands | Academia | Scenario analysis |
| Wojciech Solarz | Poland | Public administration | Policy and management |
| Lluís Brotons | Spain | Academia | Global change |
| Rubén Bernardo | Spain | Academia | Invasion ecology |
| Belinda Gallardo | Spain | Academia | Invasion ecology |
| Cristian Pérez-Granados* | Spain | Academia | Invasion ecology |
| Núria Roura-Pascual* | Spain | Academia | Invasion ecology |
| Montserrat Vilà | Spain | Academia | Invasion ecology |
| Teresa Pastor | Spain | NGO/Interest group | Policy and management |
| Garry D. Peterson* | Sweden | Academia | Global change |
| Sven Bacher | Switzerland | Academia | Invasion ecology |
| Marcus Hall | Switzerland | Academia | Environmental history |
| David Aldridge | United Kingdom | Academia | Invasion ecology |
| Guillaume Latombe | United Kingdom | Academia | Invasion ecology |
| Peter Robertson | United Kingdom | Academia | Invasion ecology |
| Helen E. Roy | United Kingdom | Academia | Invasion ecology |
| Pablo González-Moreno | United Kingdom | NGO/Interest group | Invasion ecology |

**WebPanel 1.** Step-wise participatory process to develop the management strategy based on four different scenarios for invasive alien species (IAS) in Europe. The process included 35 experts and consisted of two workshops divided into four main sessions (summarized in Figure 1).

### Part 1 (April 2020)

#### SESSION 1

***Step 1*** *– Formulation of visions for future IAS management in Europe*

The workshop participants were divided into four 'strategy breakout groups' with different stakeholders (academic and non-academic, e.g. researchers, policymakers, managers) as evenly distributed among the groups as possible. Each group separately identified and formulated different objectives (herein named visions) for the management of IAS in Europe in the coming decades until 2050.

- Output: List of management visions.

***Step 2 –*** *Selecting four future management visions of highest importance for the participants*

The management visions formulated by the different breakout groups were collected by the facilitation team and screen-shared with all participants. All participants discussed the meaning of each vision and merged those visions considered to be similar. A plenary voting decided which four visions were considered of highest importance by the participants. Each participant had four votes.

- Output: Four selected IAS management visions.

***Step 3 –*** *Development of a strategy for future IAS management in Europe*

Each of the four most voted visions was assigned to a strategy breakout group, which then developed a preliminary management strategy to reach this vision. To do so, the backcasting method was utilized, which is a way to develop plans starting from the desired future and working backwards in order to connect that future vision to the present. The core of a backcasting exercise is to determine the sequence of actions, considered in reversed order, i.e. from the end (in our case, the year 2050) to the start (present situation, year 2020), that must be taken to attain a certain goal (Robinson 1990). The advantage of backcasting as opposed to planning based on forecasting lies in the fact that reasoning from the present might limit one's ideas about the possible scope of action, resulting in steps "that are merely a continuation of present methods extrapolated into the future" (Holmberg and Robert 2000). To facilitate the formulation of their strategies, the breakout groups had access to background information from Robertson *et al.* (2020) who provide a framework about the array of actions available for IAS management and a standardized terminology.

While formulating the strategies for each individual vision, it became evident that three of them were sub-components of the fourth more overarching vision. Thus, the four partial management strategies were subsequently consolidated into one overall IAS management strategy ('*beta* version'). Therein, the suggested actions were grouped into goals that focused on specific topics. At the same time, goals were classified into four different categories: Policy, Research, Public awareness, and Biosecurity. In summary, categories correspond to broad categories of goals, goals correspond to specific management objectives identified as important by the workshop participants, and actions represent the steps required to reach these specific objectives.

- Output: Merged overall IAS management strategy ('*beta* version').

#### SESSION 2

***Step 4 –*** *Reframing & downscaling future global invasion scenarios to the European level*

For the subsequent stress-testing of the management strategy under different future conditions (see Step 6), four of the global scenarios for biological invasions (Roura-Pascual *et al.* 2021) were reframed and downscaled to the European level. To this end, the strategy breakout groups from the previous steps were reshuffled to create four new ‘scenario breakout groups’ that each comprised members from all strategy groups, ideally also representing different stakeholders. Each scenario breakout group selected one global scenario from each of the four clusters identified by Roura-Pascual *et al.* (2021) to form the basis for a downscaled European scenario for biological invasions. These clusters correspond to sets of scenarios that were grouped based on their similarities in describing potential future trajectories of biological invasions. Consequently, the four global scenarios selected for the downscaling covered widely differing future contexts, each with its own challenges and opportunities. The finally selected global scenarios, ordered from a higher to a lower level of IAS invasions, were: *Ruderal world* (cluster A), *Globalized corporation society* (cluster B), *Supernatural world* (cluster C), and*Hipster/Techno Society* (cluster D) (see Roura-Pascual *et al.* (2021)) for a more detailed description of each scenario).

For each global scenario, the respective scenario breakout group discussed elements that needed to be changed, removed or added to build a coherent picture for a European-level scenario. To structure this discussion, broad contextual developments in the following categories were considered: politics and institutions; socio-economic aspects and demographics; culture, norms and values; science and technology; and ecology and natural resources. Additionally, IAS dynamics were explicitly addressed. Changes to the original logic of the global scenario were allowed, except when they resulted in narratives that were less challenging (e.g. for future IAS management). Whenever possible, the scenario breakout groups also tried to indicate when or in which sequence certain developments occur in their scenario (timeline: present, considered as year 2020 – 2050).

In summary, the new European scenarios for biological invasions describe future framing conditions regarding social, political, economic and environmental developments in Europe (from present, year 2020, to 2050), including considerations on the status and impact of IAS. The names of the resultant scenarios were: *Lost in Europe* (derived from cluster A), *Big tech rules Europe* (cluster B), *Green local governance* (cluster C), and *Technological pseudo-panacea* (cluster D), respectively (Figure 3). A more detailed description of the European scenarios and a comparison with the global scenarios and other existing scenarios created by the climate change community (Shared Socioeconomic Pathways) will be provided in another publication.

- Output: Four European scenarios (written scenario storylines) covering from present, year 2020, to 2050.

### Part 2 (September/October 2020)

#### SESSION 3

***Step 5 –*** *Revisiting and fine-tuning the scenarios*

The four scenario breakout groups from the previous session reconvened in September/October 2020. First, they refamiliarized themselves with the European scenarios for biological invasions they developed, further refined it if additional ideas had come up in the meantime, and critically reflected on the usefulness of the scenario as a tool for stress-testing the management strategy. Particularly, it was discussed whether the scenario was challenging enough and differed enough from business-as-usual or baseline projections. At the end of this step, each breakout group had to reach a consensus on the scenario and a collective feeling about what it is like to be living under such scenario.

- Output: Four fine-tuned European scenarios for biological invasions.

***Step 6 –*** *Reviewing and stress-testing the overall IAS management strategy*

The four scenario breakout groups briefly refamiliarized themselves with the overall management strategy created in Step 3. To a certain degree, changes deemed important could still be integrated at this point. Each breakout group was asked to assume the perspective of the European scenario that they had developed and to identify the weaknesses and strengths of the overall strategy in the context of that particular future scenario ('stress test'). In particular, each breakout group assessed the feasibility of the strategy's actions under the assumptions of the respective scenario.

For each action, the assigned feasibility level (feasible, partially feasible, not feasible) had to be complemented by a brief rationale based on the scenario's logic to justify the assessment and provide context for the improvement recommendations originating from it. For example, action 1 of Policy goal 1 (i.e. related to harmonize IAS regulations at the European and the country levels) was deemed not feasible in the context of scenario "Big Tech rules Europe" (more details in Figure 3), as large corporate practices difficult coordination between countries and favour the establishment of a deregulatory attitude. In order to make it feasible, action 1 was expanded to improve coordination through the involvement of global organizations and the establishment of guidelines of best practices that countries can gradually adhere instead of enforcing agreements.

Reasons for disagreement within the group were also recorded to help identify key uncertainties. The overall aim of this step was to provide recommendations from the perspective of each European scenarios to improve the feasibility of individual management steps and the preparedness of the strategy as a whole to future challenges.

- Output: Feasibility assessments and four sets of improvement recommendations from the perspective of the four European scenarios regarding all categories of the overall management strategy.

SESSION 4

***Step 7 –*** *Integrating scenario-guided recommendations into the management strategy*

After sharing key insights from the previous step regarding challenges and recommendations from the perspective of each scenario in a brief plenary session, the former strategy breakout groups (formed in step 1) were re-established (with only minor changes in participants). By re-uniting the members of the original strategy breakout groups, these groups now combined members with detailed knowledge about all four European scenarios and about the improvement recommendations that emerged from the stress test. Each strategy breakout group was given charge of one of the four goal categories (Policy, Research, Public awareness, Biosecurity; the same categories that were developed in Step 3). Taking advantage of their combined knowledge, each strategy breakout group discussed and implemented the best way of integrating the recommendations into the respective goal category. Finally, combining the revised categories and performing some post-workshop fine-tuning (incl. coherence checks) resulted in the revised and optimized version of the overall IAS management strategy ('version 1.0').

- Output: Stress-tested and final overall strategy for future IAS management in Europe.

**WebTable 2.** Future feasibility of goals and actions of the management strategy under the assumptions of the four scenarios of biological invasions in Europe developed in the study: *Lost in Europe* (S1), *Big Tech rules Europe* (S2), *Green local governance* (S3), and *Technological (Pseudo-)Panacea* (S4) (Figure 3). The feasibility of actions was assessed qualitatively by experts, while the feasibility of goals was calculated after converting these qualitative assessments into values (Feasible: 1; Partially feasible: 0,5; Not feasible: 0) and calculating the mean. Cell colours indicate the degree of feasibility of the goals: **Feasible (1)**, **Partially feasible (0,5)** and **Not feasible (0)**.

| Goal | Action | S1 | S2 | S3 | S4 |
|---|---|---|---|---|---|
| P1 | Mean | **0.2** | **0** | **0.5** | **1** |
| | P1.1 | 0 | 0 | 0.5 | 1 |
| | P1.2 | 0 | 0 | 0.5 | 1 |
| | P1.3 | 0.5 | 0 | 0.5 | 1 |
| P2 | Mean | **0.3** | **1** | **0.3** | **1** |
| | P2.1 | 0.5 | 1 | 0.5 | 1 |
| | P2.2 | 0.5 | 1 | 0.5 | 1 |
| | P2.3 | 0 | 1 | 0 | 1 |
| P3 | Mean | **0** | **0.5** | **0.5** | **1** |
| | P3.1 | 0 | 0.5 | 0.5 | 1 |
| | P3.2 | 0 | 0.5 | 0.5 | 1 |
| | P3.3 | 0 | 0.5 | 0.5 | 1 |
| P4 | Mean | **0** | **0.5** | **0.5** | **1** |
| | P4.1 | 0 | 0.5 | 0.5 | 1 |
| | P4.2 | 0 | 0.5 | 0.5 | 1 |
| | P4.3 | 0 | 0.5 | 0.5 | 1 |
| P5 | Mean | **0** | **1** | **0.5** | **1** |
| | P5.1 | 0 | 1 | 0.5 | 1 |
| | P5.2 | 0 | 1 | 0.5 | 1 |
| P6 | Mean | **0.5** | **1** | **1** | **1** |
| | P6.1 | 0.5 | 1 | 1 | 1 |
| | P6.2 | 0.5 | 1 | 1 | 1 |
| R1 | Mean | **0.5** | **0.8** | **0.6** | **0.9** |
| | R1.1 | 0.5 | 1 | 1 | 1 |
| | R1.2 | 0.5 | 0.5 | 0.5 | 1 |
| | R1.3 | 0.5 | 0.5 | 0.5 | 0.5 |
| | R1.4 | 0.5 | 1 | 0.5 | 1 |
| R2 | Mean | **0.5** | **0.7** | **0.3** | **1** |
| | R2.1 | 0.5 | 1 | 0 | 1 |
| | R2.2 | 0.5 | 1 | 0 | 1 |
| | R2.3 | 0.5 | 0.5 | 0.5 | 1 |
| | R2.4 | 0.5 | 0.5 | 0.5 | 1 |
| | R2.5 | 0.5 | 0.5 | 0.5 | 1 |
| R3 | Mean | **0.5** | **0.7** | **0.7** | **1** |
| | R3.1 | 0.5 | 1 | 1 | 1 |
| | R3.2 | 0.5 | 1 | 1 | 1 |
| | R3.3 | 0.5 | 1 | 0.5 | 1 |
| | R3.4 | 0.5 | 0.5 | 0.5 | 1 |
| | R3.5 | 0.5 | 0.5 | 0.5 | 1 |
| | R3.6 | 0.5 | 0 | 0.5 | 1 |
| R4 | Mean | **0.5** | **1** | **0.3** | **1** |
| | R4.1 | 0.5 | 1 | 0.5 | 1 |
| | R4.2 | 0.5 | 1 | 0 | 1 |
| | R4.3 | 0.5 | 1 | 0.5 | 1 |

| Goal | Action | S1 | S2 | S3 | S4 |
|---|---|---|---|---|---|
| A1 | Mean | **0** | **0.1** | **0.5** | **0.9** |
| | A1.1 | 0 | 0 | 0.5 | 1 |
| | A1.2 | 0 | 0 | 0.5 | 1 |
| | A1.3 | 0 | 0 | 0 | 0.5 |
| | A1.4 | 0 | 0.5 | 1 | 1 |
| A2 | Mean | **0.5** | **0.5** | **0.5** | **1** |
| | A2.1 | 0.5 | 1 | 0.5 | 1 |
| | A2.2 | 0.5 | 0 | 0.5 | 1 |
| A3 | Mean | **0.5** | **0.6** | **0.8** | **0.8** |
| | A3.1 | 0.5 | 0.5 | 1 | 0.5 |
| | A3.2 | 0.5 | 1 | 0 | 1 |
| | A3.3 | 0 | 0.5 | 1 | 1 |
| | A3.4 | 1 | 0.5 | 1 | 0.5 |
| B1 | Mean | **0.2** | **0.2** | **0.3** | **0.8** |
| | B1.1 | 0.5 | 0.5 | 1 | 1 |
| | B1.2 | 0 | 0 | 0 | 1 |
| | B1.3 | 0 | 0 | 0 | 0.5 |
| B2 | Mean | **0.5** | **0.3** | **0.2** | **1** |
| | B2.1 | 0 | 0 | 0 | 1 |
| | B2.2 | 1 | 0.5 | 0.5 | 1 |
| | B2.3 | 0.5 | 0.5 | 0 | 1 |
| B3 | Mean | **0.6** | **0.3** | **0.6** | **0.9** |
| | B3.1 | 0.5 | 0.5 | 0 | 1 |
| | B3.2 | 0.5 | 0.5 | 1 | 1 |
| | B3.3 | 0.5 | 0 | 0.5 | 1 |
| | B3.4 | 1 | 0.5 | 1 | 0.5 |
| | B3.5 | 0.5 | 0 | 0.5 | 1 |
| B4 | Mean | **0.5** | **0.5** | **0.4** | **1** |
| | B4.1 | 0.5 | 0.5 | 0 | 1 |
| | B4.2 | 0.5 | 0.5 | 0.5 | 1 |
| | B4.3 | 0.5 | 1 | 0.5 | 1 |
| | B4.4 | 0.5 | 1 | 0.5 | 1 |
| | B4.5 | 0.5 | 0 | 0.5 | 1 |
| | B4.6 | 0.5 | 0 | 0.5 | 1 |
| B5 | Mean | **0.6** | **0.5** | **0.5** | **0.9** |
| | B5.1 | 1 | 0.5 | 0.5 | 1 |
| | B5.2 | 0.5 | 0.5 | 0.5 | 1 |
| | B5.3 | 0.5 | 0.5 | 0.5 | 1 |
| | B5.4 | 0.5 | 0.5 | 0.5 | 0.5 |
| | B5.5 | 0.5 | 0.5 | 0.5 | 1 |
| B6 | Mean | **0.5** | **0.2** | **0.7** | **1** |
| | B6.1 | 0.5 | 0.5 | 0.5 | 1 |
| | B6.2 | 0.5 | 0 | 1 | 1 |
| | B6.3 | 0.5 | 0 | 0.5 | 1 |
