## Supplementary Material (WebPanel 2) for "A scenario-guided strategy for the future management of biological invasions"

### IAS MANAGEMENT STRATEGY

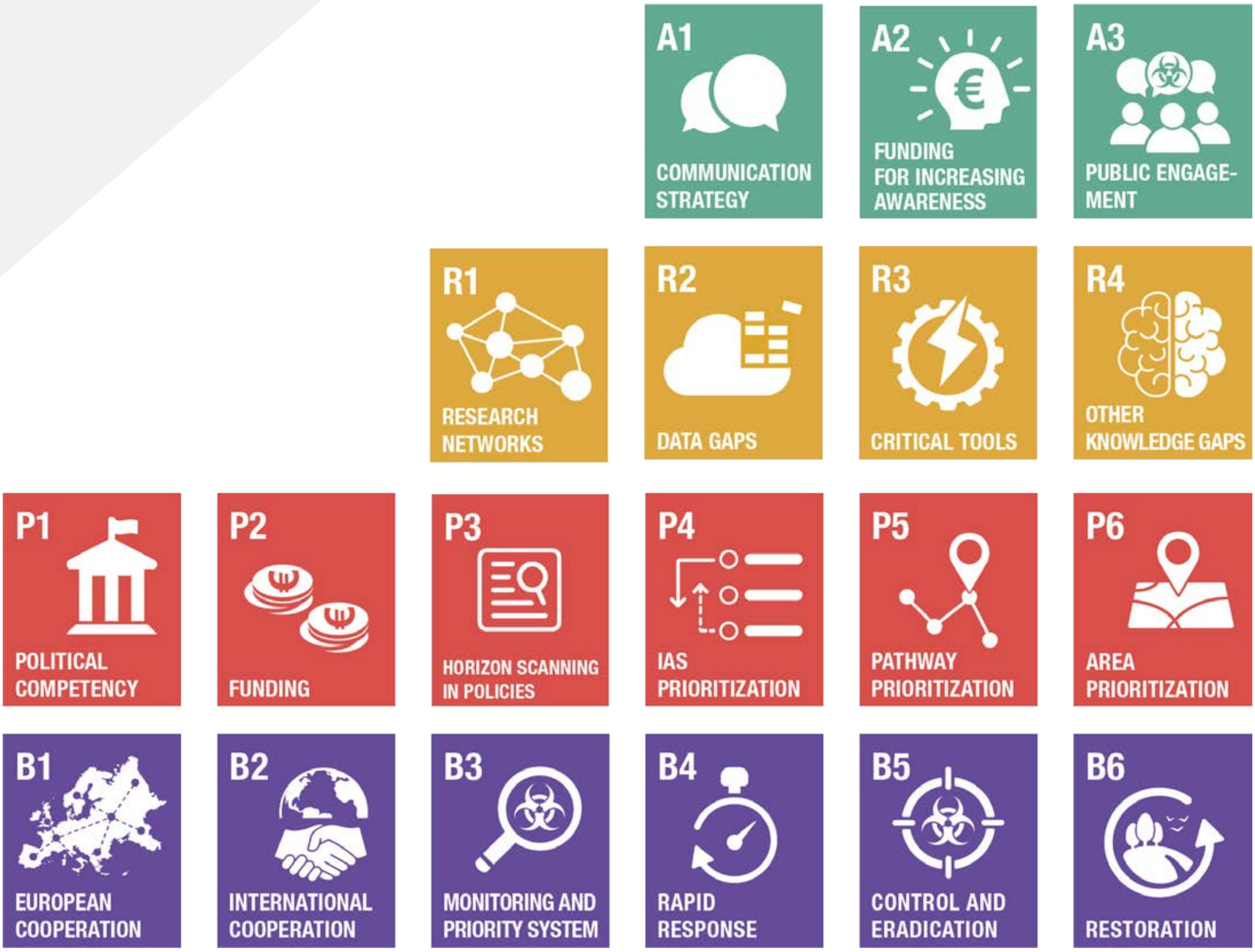

### IAS Management Strategy

**Outline** ........ 2
Preamble ........ 3
Overall vision of the strategy ........ 3
IAS management strategy ........ 4

**Policy**
Goal P1: Political competency ........ 4
Goal P2: Funding ........ 5
Goal P3: Horizon scanning in policies ........ 7
Goal P4: IAS prioritization ........ 8
Goal P5: Pathway prioritization ........ 10
Goal P6: Area prioritization ........ 11

**Research (data / tools / information)**
Goal R1: research networks ........ 12
Goal R2: Data gaps ........ 13
Goal R3: Critical tools ........ 14
Goal R4: Other knowledge gaps ........ 16

**Public awareness and communication with stakeholders**
Goal A1: Communication strategy ........ 17
Goal A2: Funding for increasing awareness ........ 18
Goal A3: Public engagement ........ 19

**Biosecurity, monitoring, management and restoration**
Goal B1: European biosecurity regime ........ 20
Goal B2: International cooperation ........ 22
Goal B3: Monitoring and assessment system ........ 23
Goal B4: Rapid response ........ 24
Goal B5: Control and eradication ........ 25
Goal B6: Restoration ........ 26

References ........ 27

### PREAMBLE

The IAS management strategy presented here was developed in 2020 during a participatory workshop process together with stakeholders. We define stakeholders following Shackleton *et al.* (2019) as "any individual, group or organisation who is affected (positively or negatively) by invasive species, or who has the capacity to promote or limit the spread of invasive species [...]. Stakeholders include the public/citizens (affected by and/or responsible for the spread and/or control of invasive species), researchers, government departments (responsible for the management of invaded areas or as policy makers), non-governmental organisations (NGOs), businesses and industry, and many other groups [...]".

The management strategy aims to encompass management actions for the whole of Europe, i.e. Member States of the European Union (EU) as well as European states not currently members of the EU (as of 2020).

The strategy is aiming to reach its overall vision (see below) until the year 2050, i.e. within a period of 30 years since 2020. Although goals are also stated with this general time horizon in mind, some of them can be seen as stepping stones towards the overall vision and their achievement would be desirable for earlier than 2050.

### OVERALL VISION OF THE STRATEGY

*By 2050, the harmful impacts of invasive alien species (IAS) in Europe (EU-Member States and non-EU states) are substantially reduced compared to today.*

More specifically, by 2050 no or few IAS with potentially harmful impacts (i.e. negative effects on biodiversity, socio-economics or human health) are being introduced, establish or spread in Europe. The harmful impacts of IAS are low in all types of habitats and do not increase over time. Management of IAS can be done with minimised with non-target effects and can be performed cost-effectively. ['Management' is here defined in a broad sense, following the framework by Robertson *et al.* (2020), i.e. including prevention, captive management, rapid eradication and long-term management.] There is high public awareness of the harmful impacts of some IAS, and people (individuals or relevant stakeholders, such as gardening/forestry/pet market enterprises, etc.) act responsibly to further avoid the introduction, establishment and spread of harmful IAS.

---

In contrast, the current situation is as follows:

- Many new IAS are arriving, establishing and spreading each year - we are in a leaking boat (Seebens *et al.* 2017; Pyšek *et al.* 2020).
- The free market within the EU (i.e. free movement of goods) counteracts effective invasion management and the enforcement of regulations.
- There is a lack of knowledge about the impacts of most IAS. This is even true for well investigated taxonomic groups such as birds (Evans *et al.* 2016, 2020).
- IAS management is conducted on a limited number of sites which were not selected as a result of evidence-based priorities (Dana *et al.* 2014).
- There is low public awareness about the harmfulness of IAS (Courchamp *et al.* 2017).

### IAS MANAGEMENT STRATEGY

To reach the overall vision, we have formulated goals that are grouped into the following four categories: policy, research, public awareness, and biosecurity. As such, the responsibility for implementing the goals lies with different levels of political organization as well as public and private institutions, as seen appropriate, which are required to collaborate and coordinate their actions.

#### Policy

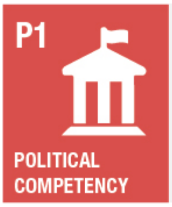


##### Goal P1:
Strengthened IAS competency of political agencies in Europe

*GOAL: By 2050, European political agencies have high competency and effectiveness in preventing and mitigating harmful impacts of IAS.*

**Actions to reach this goal:**

1. Harmonize IAS regulations both at the European and the country level so that management actions can be better coordinated.

    - Ensure, as a minimum, that all European countries (incl. non-EU states) have IAS regulations and transparently communicate them both at the European and the country level to all involved political agencies.
    - Invite European and global international organizations to participate in or lead the harmonization, providing recommendations for policy development and guidelines of best practice of management, enabling their iterative refinement (e.g. for considering regional/local particularities), monitoring their implementation (with or without enforcement), and facilitating the communication / coordination between EU Member States and non-EU states.
    - If full harmonization of IAS regulations is not possible, and depending on the level of cooperation between European countries,
        - encourage countries to choose from approaches included in the guidelines of best practice that best suit their management situation (“participatory mode of functioning”);
        - gradually increase harmonization of the IAS regulations by e.g. including knowledge transfer to learn from each other’s experiences, shared minimum management principles/actions, joint development of new management options, or coordination of management.

2. Strengthen the legislative framework at the European (incl. non-EU states) and country level, so that management actions are more effective.

    - Agree upon high-level principles of IAS legislation and management outcomes to be achieved by all European countries, but not aiming at full agreement on finer details across Europe, e.g. species listing.
    - If the level of cooperation between European countries allows, a European agency or intergovernmental agreement for IAS management shall be responsible for the communication and coordination between European countries, the provisioning of recommendations and guidelines (incl. due consideration of regional/local particularities), and the liaison to global organizations and conventions.
    - On a worldwide level, promote the establishment of a global IAS organization/platform, e.g. within the UN.

3. Increase the effectiveness of political agencies and the legislative framework by actively involving members of the private sector that have an interest in the successful management of economically important IAS.

---

**Current situation:**

- EU IAS Regulation (1143/2014).
- Other regulations at the EU level or country level cover other aspects, e.g. plant health or IAS affecting human or animal health, ballast water convention.
- Lack of or inefficient collaboration/communication between EU and non-EU countries.
- Existence of a number of bodies that assist the European Commission in the implementation of the IAS Regulation at different levels: Committee on IAS, Invasive Alien Species Expert Group (IASEG), Scientific Forum on IAS, the Working Group on IAS, and Bern Convention Expert Group on IAS, among others. For more information, see the website of the European Commission on IAS.

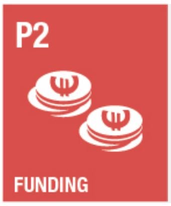


#### Goal P2:
Funding (for research, monitoring, management, restoration)

*GOAL: By 2050, there is funding to fill critical research gaps and maintain the monitoring/management and restoration activities related to IAS at the European (incl. non-EU members) and country level.*

**Actions to reach this goal:**

1. Identify funding needs and resources (public and private) at the European and country level.

    - Invite European and global international organizations to lead this process, if possible forming an expert panel and providing recommendations for prioritizing the funding needs, while considering country-specific or regional differences.

- Support the search for funding sources with a communication campaign about IAS impacts and their management to increase awareness in the general public and in potential public and private funders.
- If the level of cooperation between European countries allows, a European agency or intergovernmental agreement for IAS management shall be responsible for identifying funding needs (in collaboration with above-mentioned expert panel) as well as the collection of funds and their fair and transparent distribution between all European countries.

2. Establish and/or revise public funding priorities based on action 1.

- Enhance the coordination between countries within and beyond Europe based on existing (e.g. CBD, Bern Convention, Ballast Water Management Convention) and new agreements.
- Consider the short- and long-term costs of IAS including the necessity of strong preventive biosecurity as well as of long-term management for already established IAS.

3. Secure long-term funding at the European and country level.

- Seek maximum independence of funding from legislative periods.
- Take advantage of cost-reducing synergy effects by targeting high-impact 'umbrella' IAS for which the management is of interest to (economic) sectors with high funding capacities (e.g. industry) and is at the same time in line with management objectives for other IAS.
- Create concrete (economic) incentives for the private sector to contribute funds to IAS management. To engage key economic sectors, clearly demonstrate cross-sectoral impacts of IAS that go beyond negative consequences for biodiversity conservation.
- Increase efforts to involve volunteer work to free up available funding, which can then be used for other purposes than personnel.
- If the level of public awareness and cooperation allows, consider introducing a tax on donor activities that are susceptible to introducing alien species ('polluter-pays principle', following Directive 2004/35/EC on environmental liability with regard to the prevention and remedying of environmental damage).

---

**Current situation:**

- General lack of funding for research, monitoring, management and restoration, with strong differences among (and partly within) European countries.
- Particular and critical lack of funding for long-term initiatives.
- The European Commission is supporting actions on IAS through existing financing instruments (LIFE, Horizon 2020, Horizon Europe, rural development programmes, etc), but there is not a specific funding programme dedicated to IAS (such as Working for Water in South Africa, etc.).

P3

HORIZON SCANNING IN POLICIES

#### Goal P3:

Incorporating Horizon Scanning recommendations in IAS policies

*GOAL: By 2050, Horizon Scans are regularly carried out at the European (incl. non-EU members) and country level, and their results are incorporated in IAS policy.*

**Actions to reach this goal:**

1. Agree on a harmonized horizon scan (HS) protocol for alien species at the European level (incl. non-EU states). [Horizon scanning is defined as a systematic examination of potential threats and opportunities of multiple species within a given context to prioritize or rank species (Roy *et al.* 2014)]

   - Invite European and global international organizations to lead this action, if possible forming an expert panel. Include local expertise whenever possible.
   - Connect HS to mechanisms which ensure that it effectively triggers further action, e.g. prioritization of management actions within a risk management framework.
   - If a low level of cooperation prevents agreement, ensure as a minimum that HS protocols exist for each individual European country according to their specific perspectives, which will facilitate the subsequent adoption of a harmonized protocol.
   - For cases where capacities to carry out HS are limited, develop guidelines for focusing HS on priority issues (focusing on e.g. IAS with highest impacts, management feasibility, or threat to multiple sectors of high interest such as industry, agriculture, water, human health).
   - Make the protocol available in different languages within an online system that is easy to use and allows to conduct HS with reduced needs for travelling and coordination between collaborators.

2. Regularly carry out HS at the European (incl. non-EU members) and country level. Europe-wide HS helps identify IAS for common action, while country-level HS helps identify IAS of particular interest for a given country.

   - Invite European and global international organizations to lead and coordinate this action (see also action 1).
   - If possible, implement flexible evidence- and knowledge-based update intervals instead of fixed update intervals such as “every 5-10 years”.
   - If possible, integrate and continuously update HS data in a centralized European data platform, which ideally also allows to conduct HS online (see action 1). Otherwise, as a minimum, build a simple, standardized online interface that facilitates regular and consistent revisions.

3. Incorporate HS recommendations in European IAS policy regarding IAS management at all invasion stages, or else in national policies at least.

    - Support this process by strengthening relationships between and visibility of all (public and non-public) stakeholders, and by identifying influential individuals/groups (incl. grassroot movements) that can push for changes in European and/or national IAS policy.
    - If the level of cooperation among European countries as well as between Europe and the rest of the world allows, strengthen the competences of global conventions (and the organisations curating them) so that such policy can be enforced onto national governments.

---

**Current situation:**

- According to EU IAS Regulation (1143/2014), Member States should have implemented functional structures and a surveillance system of IAS included in the EU list but HS is not considered in the regulation. The current list used the previous existing HS exercise.
- Horizon Scanning is currently done at the EU-level (for terrestrial and recently for marine environments) and in few non-EU states (Roy *et al.* 2019; Tsiamis *et al.* 2020).

#### Goal P4:

Regular revision of IAS priority list

*GOAL: By 2050, IAS priority lists at the European (incl. non-EU states) and country level are legally implemented and regularly revised, and "priority" is assessed based on different types of impacts, including those on biodiversity, socio-economics and human health.*

**Actions to reach this goal:**

1. Identify and prioritize IAS for their management at the European (incl. non-EU states) and country level.

    - Use updated standard risk or impact assessments (e.g. S/EICAT; Blackburn *et al.* 2014; Hawkins *et al.* 2015; Bacher *et al.* 2018; IUCN 2020) and transparent prioritization rationales (give particular consideration to management feasibility of a species and its impact on key economic sectors) (Booy *et al.* 2017, 2020; Adriaens *et al.* 2019).
    - Develop national and/or (eco-)regional[1] IAS priority lists in parallel to a European IAS priority list since they may reflect important, more localized particularities of IAS management. Regions can be defined based on a variety of concepts, e.g. administrative subunits of a country, areas with the same language or cultural legacy. Especially in the context of IAS management, ecological parameters can be used to define eco-regions.

[1] The "eco"-regional approach can make sense since a region can be defined in different terms, an important one for the management topic being ecological parameters (instead of e.g. administrative sub- or super-units of a country or areas with the same language, cultural legacy, etc.)

- Solicit suggestions from European and global international organizations for the European IAS priority list, which consider pre-existing European IAS legislation (e.g. the 'Union list' in EU IAS Regulation 1143/2014) and include concrete recommendations for harmonization between countries (considering also regional networks, e.g. Scandinavia, BENELUX).
- Ensure the implementation of the IAS priority lists in respective European IAS legislation.

2. Regularly revise European, regional and national IAS priority lists.

   - Make lists dynamic and flexible, e.g. allowing the removal of species for which no longer any prospect of effective management exists, or emergency protocols for swift inclusion of newly identified high-risk IAS.
   - If possible, implement flexible evidence- and knowledge-based update intervals instead of fixed update intervals such as "every 5-10 years".
   - If possible, integrate and continuously update data related to the IAS priority listing in a centralized European data platform. Otherwise, as a minimum, build a simple, standardized online interface that facilitates regular and consistent revisions.
   - If the level of cooperation between European countries allows, a European agency or intergovernmental agreement for IAS management shall be responsible for facilitating and coordinating the regular revisions.

3. Incorporate different types of impacts for the prioritization of IAS, including impacts on biodiversity, socio-economics and human health.

   - Use the most recent findings from multidisciplinary targeted research (e.g. ecology, economics, sociology) across different countries (see section Research).
   - Whenever possible, quantify impacts not exclusively in monetary terms; still, the economic impact should be quantified especially for (societal) settings in which knowledge about financial benefits or losses is the only effective incentive for management.
   - Encourage the input from global organizations from different sectors (e.g. biodiversity, plant protection, WHO, WTO).
   - Weight the different impacts in a transparent way (e.g. implementing co-management boards, multi-criteria decision analyses considering points of view of different stakeholders) to ensure appropriateness and acceptance of prioritization.

---

**Current situation:**

- An EU IAS priority list is regularly revised according to the EU IAS Regulation (1143/2014), while there is no IAS priority list at the level of the whole European continent.
- EU Member States are able to apply for including new species in EU IAS regulation (1143/2014).
- EU IAS Regulation (1143/2014) has prioritized IAS based on their biodiversity impacts, with other impacts considered as aggravating factors.
- The current list of IAS of EU concern is relatively short and of limited usefulness to guide the management of IAS across the EU.
- There is no white-list approach in the EU.

#### Goal P5:
Regular revision of pathway priority list

*GOAL: By 2050, European countries have identified pathways of introduction and spread, and agreed on a list of priority pathways that is legally implemented and regularly revised.*

**Actions to reach this goal:**

1. Identify and prioritize pathways of introduction and spread for IAS management at the European (incl. non-EU states) and country level.

    - Use standard protocols (e.g. CBD standardized pathway scheme; CBD 2014; Harrower *et al.* 2020), metrics that facilitate comparisons across countries, and transparent prioritization rationales. Give particular attention to the identity and number of IAS per pathway, impact on key economic sectors, as well as the identification and involvement of specific actors that are contributing to or suffering from the importance of certain pathways due to their socio-economic activities.
    - Develop national and/or (eco-)regional pathway priority lists in parallel to a European pathway priority list since they may reflect important, more localized particularities of IAS management.
    - Solicit suggestions from European and global international organizations for the European pathway priority list, which consider pre-existing European IAS legislation (e.g. pathway prioritization within EU IAS Regulation 1143/2014) and include concrete recommendations for harmonization between countries (considering also regional networks, e.g. Scandinavia, BENELUX).
    - Ensure the implementation of the pathway priority lists in respective European IAS legislation.

2. Regularly revise the European, regional and national pathway priority lists.

    - If possible, implement flexible evidence- and knowledge-based update intervals instead of fixed update intervals such as “every 5-10 years”.
    - If possible, integrate and continuously update data related to the pathway priority listing in a centralized European data platform. Otherwise, as a minimum, build a simple, standardized online interface that facilitates regular and consistent revisions.
    - If the level of cooperation between European countries allows, a European agency or intergovernmental agreement for IAS management shall be responsible for facilitating and coordinating the regular revisions.

---

**Current situation:**

- According to EU IAS Regulation (1143/2014), member states should have identified the pathways of unintentional introduction and spread of invasive alien species of Union Concern.

- Differentiating between pathways of primary introduction (e.g. intercontinental introductions to major ports) and of subsequent secondary introduction (e.g. within country transport to smaller towns or natural spread of introduced species) is still difficult but would help to use limited management resources more efficiently.

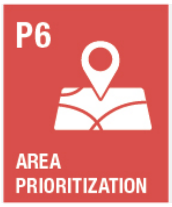


#### Goal P6:

Regular revision of priority areas for IAS management

*GOAL: By 2050, European countries have elaborated a list of priority areas for IAS management that is legally implemented and regularly revised.*

**Actions to reach this goal:**

1. Identify and prioritize areas for IAS management (as suggested by McGeoch *et al.* (2016)) at the European and country level.

    - Use standard protocols, metrics that facilitate comparison between countries, and transparent prioritization rationales.
    - Develop national and/or (eco-)regional lists of priority areas for IAS management in parallel to a European priority list since they may reflect important, more localized particularities of IAS management.
    - Solicit suggestions from European and global international organizations for the European list of priority areas for IAS management, which consider pre-existing European IAS legislation as well as the need of neighbouring countries to establish and agree on a common set of priorities for border regions (increased communication).
    - Ensure the implementation of the priority lists in respective European IAS legislation.

2. Regularly revise the European, regional and national lists of priority areas for IAS management.

    - If possible, implement flexible evidence- and knowledge-based update intervals instead of fixed update intervals such as “every 5-10 years”.
    - If possible, integrate and continuously update data related to the priority areas in a centralized European data platform. Otherwise, as a minimum, build a simple, standardized online interface that facilitates regular and consistent revisions.
    - If the level of cooperation between European countries allows, a European agency or intergovernmental agreement for IAS management shall be responsible for facilitating and coordinating the regular revisions.

---

**Current situation:**

- In EU Regulation (1143/2014), the need to identify priority areas for IAS management is not mentioned.
- Currently no specific priority areas for IAS management across Europe (there are areas that receive a stronger protection than others, e.g. Natura 2000 sites, but these are not selected based on a specific consideration of IAS). IAS criteria are insufficiently developed in the assessment criteria for the conservation status of Natura 2000 habitats.

### Research (Data / Tools / Information)

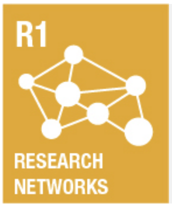


#### Goal R1:
Establishment of research networks for IAS management

*GOAL: By 2050, strong and active research networks regarding IAS management have been established at the European level and beyond.*

**Actions to reach this goal:**

1. Identify and contact relevant stakeholders related to IAS research (e.g. researchers, managers, policy-makers, public administrators, NGOs, etc.).

2. Strengthen current relevant networks and platforms of IAS researchers, managers, policy-makers and other stakeholders. This includes global organizations (e.g. IUCN, WTO, WHO, UN). Create public-private partnerships at an international level.

3. Ensure that all stakeholders recognize the need for collaborative work, clear communication and sharing data. Achieve this by following the collaborative, multisectoral and transdisciplinary "One Health" approach (FAO et al. 2019) (including incorporation of Nature's Contributions to People), providing incentives to share data and applying a variety of engagement methods (e.g. role-playing games).

4. Establish and actively maintain strong links to international initiatives, e.g. for IAS data collection and curation, and whenever possible extend these links beyond the European context.

    - Potential approaches include online fora, co-management boards or multi-criteria decision analyses (Dana et al. 2014) to incorporate a diversity of values and perspectives.
    - Special attention needs to be given to potential differences between EU members and non-EU states.

---

**Current situation:**

- The Joint Research Centre (JRC) of the European Commission developed the European Alien Species Information Network (EASIN), an online platform that aims to facilitate access to existing information on alien species.
- Lack of or inefficient collaboration and communication between European and non-European countries.
- Effective IAS management frequently needs other types of knowledge than what researchers currently investigate.
- Communication across disciplinary boundaries is often hampered by unclear communication based on jargon and the 'logical alien' effect (see Millgram 2015; Jeschke *et al.* 2019).

R2

DATA GAPS

#### Goal R2:
Targeted research on critical data gaps

*GOAL: By 2050, critical data gaps (related to e.g. the quality, quantity, storage, harmonization, accessibility, processing and updating of IAS data) are regularly identified, and targeted research is continuously closing them.*

**Actions to reach this goal:**

1. Improve existing and develop missing or complementary data standards that are straightforward to apply.

    - Follow (optimized) standardized protocols (e.g. FAIR principles, Darwin Core terms) and make data openly accessible.
    - Reach out to policy-makers in the process explaining the benefits of data standards, and invite global organizations to promote the data standards for their international, ideally worldwide, acknowledgement.

2. Establish and implement a standardized data infrastructure, allowing integration of data from various sources (e.g. earth observations, remote sensing, text mining algorithms, image recognition) and improved data flow across spatiotemporal scales.

    - Consider the adoption of technologies that were developed for other purposes.
    - If the level of cooperation between European countries allows, a European agency or intergovernmental agreement for IAS Management or global organization shall be responsible for the development and maintenance of an easy-to-use online platform to facilitate data integration across European countries.

3. Identify species for which we currently do not have sufficient data (due to actual data-deficiency or no existing alien population, e.g. Watkins et al. 2021) about their (potential) introduction pathways, distribution and impacts (see next action 4), and for which assessments of their (potential) impacts should be done. Prioritize these species based on results from horizon scanning, using open-access databases and cooperation through research networks supported by international organizations.

4. Identify potential priority areas and pathways (based on transparent prioritization rationales) for which we currently do not have sufficient data, in particular regarding native and non-native species present there and human pressures.

5. Expand and intensify the collection of data related to invasions, in particular on data-deficient IAS (e.g. species distribution, impacts) and on priority areas and pathways (see previous actions 3 and 4 above).

    - Include data that were sourced outside the invasion context but are nevertheless relevant for IAS management.

- Continuously map and update the distribution of IAS across European countries at a fine spatial scale,
  - making use of standardized mapping protocols, newest available technologies (e.g. remote sensing, DNA barcoding, environmental DNA, smart data capture for high degree of automation) and citizen science. Procure the manpower that is needed to apply these techniques (e.g. for calibrating tools/sensors, training AI algorithms);
  - encouraging (or if possible requiring) owners of IAS distribution data (governmental institutions as well as private companies) to share their data, and increasing database integration with international initiatives (e.g. CABI, GBIF);
  - integrating and continuously updating distribution data if possible in a centralized European data platform.
- Apply automated approaches to managing and analyzing big datasets, and implement incentives (e.g. funding) for open science.

---

**Current situation:**

- Several IAS databases (European and global) that are already available are an important basis from which to start; progress is being made with harmonization between them (e.g. CABI ISC, DAISIE, EASIN, IUCN GISD).
- Certain IAS databases are not openly available (but there is progress).
- Certain IAS databases are outdated/not kept up to date anymore (e.g. DAISIE, GRIIS).
- Interoperability between databases is often extremely complex or non-existent (e.g. taxonomy).
- Collation of data from different IAS databases for more statistical power/ synergy effects is still difficult.
- Citizen science for IAS in on the rise in Europe with many initiatives emerging in recent years, but there is a lack of guidance, harmonization, openness of data and data interoperability.
- Making data and research findings available with open access is slowly gaining more attention but is still not widespread enough for facilitating collaborative research and management.
- Taxonomic expertise in general and related to IAS in particular is dwindling or completely missing.

R3 CRITICAL TOOLS

#### Goal R3:

##### Targeted research on critical tools related to IAS impact/risk assessment and management

*GOAL: By 2050, critical gaps related to tools for IAS impact/risk assessment (incl. for horizon scans) and their management are regularly identified and targeted research is continuously closing them.*

**Actions to reach this goal:**

1. Improve existing impact assessment standards (e.g. S/EICAT; Blackburn *et al.* 2014; Hawkins *et al.* 2015; Bacher *et al.* 2018; IUCN 2020) and develop missing or complementary standards (e.g. for beneficial impacts), considering all impact dimensions (including impacts on biodiversity, socio-economics and human health) and knowledge from local people and practitioners.

2. Improve existing standards for assessing IAS pathways (e.g. CBD scheme based on Hulme *et al.* (2008)); see also CBD (2014) and Harrower *et al.* (2020)) and their management.

3. Develop standards for assessing the vulnerability and management of priority areas (cf. Roura-Pascual 2009).

4. Develop an evidence-based and adaptive approach to guide management decisions. Review and analyze the effectiveness of management actions that have already been implemented, in particular by assessing differences in effectiveness across techniques, regions and taxa.

5. Develop novel management techniques (e.g. drones and robots that automatically detect or eradicate IAS, internet-based detection of alien species occurrences, eDNA; Martinez *et al.* 2020).

    - Catalyze and incentivize technology innovation and dual-use technology (i.e. technologies originally developed for military, intelligence, human health or other purposes).
    - Attract experts in novel relevant technologies to join the networks of researchers and other stakeholders.

6. Ensure full adoption, integration and implementation of tools at the European and country level and, if applicable, by international organisations.

    - Create, advance, and maintain the regulatory frameworks necessary for technology development and application.
    - Follow (optimized) standardized protocols (e.g. FAIR principles, Darwin Core terms) and make tools openly accessible.
    - Apply modular protocols and workflows that allow the dynamic integration of new information by any stakeholder (cf. Vanderhoeven *et al.* 2017).
    - Establish an active program for identifying novel and effective tools developed within and beyond Europe.
    - Consider incentives (e.g. funding) to foster open science.

---

**Current situation:**

- According to EU IAS Regulation (1143/2014), Member States should have implemented functional structures and a surveillance system of IAS included in the EU list.
- Lack of robust and widely used, standardised methodologies for impact quantification, cost-benefit analyses, simulations of management effectiveness (e.g. for identifying effectiveness thresholds), development of standards.
- It is unclear whether we are using the most appropriate indicators and metrics for IAS and their impacts.

R4
OTHER KNOWLEDGE GAPS

#### Goal R4:
Targeted research on other critical knowledge gaps

*GOAL: By 2050, other critical knowledge gaps regarding our understanding of IAS and their management are regularly identified, and targeted research is continuously closing them.*

**Actions to reach this goal:**

1. Establish and implement a standardized terminology toolbox, allowing integration of various knowledge systems (including indigenous and local knowledge) and data sources (e.g. earth observations, remote sensing, horizon scanning protocols, risk assessment protocols, text mining algorithms, image recognition). Flexibility should be considered in standardized terminology, since terms can become obsolete and are not used consistently by different sectors. This can be achieved by ontologies which clarify how different terms relate to each other (Madin *et al.* 2008).

2. Identify knowledge gaps and define research targets that need to be addressed.

   - Take advantage of established networks of researchers and other stakeholders. Promote multi-disciplinary collaboration of researchers, e.g. through specific funding calls.
   - Substantially improve understanding of e.g.:
     - interactions of IAS with climate change and other anthropogenically driven pressures on biodiversity,
     - methods for monitoring and eradicating/controlling IAS,
     - applied theory in IAS management (e.g. relationship to basic ecological concepts and hypotheses),
     - economics of IAS management,
     - restoration ecology (incl. novel ecosystems),
     - IAS impacts (incl. strategies of how to live with IAS and adapt to their impacts, or even taking advantage of their positive impacts),
     - approaches for effective science communication/outreach and teaching.

3. Regularly revise research targets and cross-cutting research areas and define new ones if necessary.

---

**Current situation:**

- Taxonomic expertise in general and related to IAS in particular is dwindling or completely missing.
- Research is typically focused on one or few biodiversity drivers, and the interactions of multiple socio-ecological drivers are largely unknown.
- Indigenous and local knowledge does not receive sufficient attention and much of it is lost.
- Making data and research findings available with open access is slowly gaining more attention but is still not widespread enough for facilitating collaborative research and management.

### Public awareness and communication with stakeholders

A1 COMMUNICATION STRATEGY

#### Goal A1:

Cross-sectoral IAS communication strategy

*GOAL: By 2050, a cross-sectoral IAS communication strategy and centralized communication platform has been developed and established, and is actively implemented at the European (incl. non-EU members) and country level to achieve high public awareness of IAS, their impacts and their management in Europe, a requirement for creating an effective legal framework.*

**Actions to reach this goal:**

1. Form a working group on IAS communication at the European level with active participation of stakeholders from all sectors (including e.g. NGOs, science journalists and key actors with control over a sector or ecosystem) and European regions.

   - Involve global networks and promote the self-motivated participation of non-EU countries through clear advantages (e.g. better coordination) and low participation costs.
   - Develop an inclusive platform for knowledge sharing and facilitate cooperation across different contexts/circumstances (e.g. pluralism of languages, issues, concerns and values - cultural values of nature vs. nature for production).

2. Develop a cross-sectoral communication strategy regarding IAS and their management (led by the communication working group; see action 1) to raise awareness and enable extensive collaboration and knowledge transfer.

   - Implement approaches for effective science communication outreach. Foster diverse types of dialogue and exchange between stakeholders (e.g. involving managers but also communities with strong links to and extended practical knowledge about nature (e.g. farmers, gardeners, hunters, birdwatchers).
   - Bring sectors together by highlighting the diversity of IAS impacts, how they are connected across sectors and space, and the benefits of addressing synergistic impacts of IAS and other environmental changes (e.g. climate change, habitat fragmentation, over-fertilization) in an integrated way (consider the role of different concepts of valuation of nature).

3. Establish and regularly update a centralized, multilingual European platform and infrastructure for knowledge transfer and raising public awareness as an integral tool of the cross-sectoral IAS communication strategy.

   - Ensure the platform is reliable, has an open data sharing policy, and follows standardized protocols (e.g. FAIR principles) so that new global, regional or national databases could build upon one another and share data.

- Encourage the continued use of the platform (including the interactive updating of data) by all stakeholder groups, including making sure validity, high quality and relevance of data by implementing a peer-review process.
- If possible, link the communication platform to a centralized European IAS data portal to increase the rate, extent and efficiency of knowledge transfer and communication.

4. Prepare a dedicated education curriculum for schools conveying the value of biodiversity and the potential consequences of biological invasions, as well as addressing the phenomenon of shifting baselines (Soga and Gaston 2018) in the perception of invasions.

    - Use teaching methods that are up-to-date and can adapt to different backgrounds of learners (e.g. pronounced technophily) and attitudes towards the topic.

---

**Current situation:**

- Lack of or inefficient collaboration/communication between European and non-European countries.
- Cross-sectoral communication is very complicated, as every sector has its own jargon and logic, is in its own silo (Millgram 2015; Jeschke *et al.* 2019).

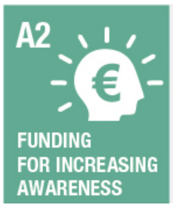


#### Goal A2:
Financial resources for raising public awareness

*GOAL: By 2050, there is dedicated funding at the European (incl. non-EU states) and country level for raising public awareness.*

**Actions to reach this goal:**

1. Inform potential funders (public and private) about the potential consequences of IAS across all sectors, the importance of their management, and the need to enlist all sectors for collaboratively developing effective solutions.

    - Highlight country- or region-specific IAS issues to amplify awareness of funders with close links to those countries or regions.

2. Secure dedicated resources (public and private) for raising public awareness by involving local, national or global institutions.

    - Consider setting up organizations with long-term funding that (i) link IAS management with direct benefits for people (e.g. regarding training and employment; see the South African 'Working for Water' programme) or (ii) bridge different sectors engaging their stakeholders to contribute towards a shared objective (see e.g. 'Water Funds').

---

**Current situation:**

- NGOs as funders do not currently strongly support IAS management in Europe due to conflicting ethical views (in particular, killing organisms is often seen as unethical, even if the species is invasive).

A3 PUBLIC ENGAGE-MENT

#### Goal A3:
Involve the general public and other relevant stakeholders in IAS management

*GOAL: By 2050, effective means are in place to involve the general public and other relevant stakeholders in IAS management in all European countries.*
*Actions to reach this objective:*

**Actions to reach this goal:**

1. Produce dissemination material for motivating the general public to act upon harmful IAS.

    - Adapt the dissemination material to people's interests and needs, considering different levels of education and ages and including innovative ways of communication (e.g. games, comics, social media).
    - Create material for specific regions or sectors, highlighting concrete invasion impacts in them or the most harmful IAS in order to engage the local public.
    - Explicitly counter misinformation and address the phenomenon of shifting baselines (Soga and Gaston 2018).

2. Produce comprehensive technical guidance on IAS management, control and eradication in Europe based on the most recent findings from targeted research, and disseminate it in various formats to relevant stakeholders.

    - Consider creating communities of practice for connecting stakeholders and promoting training activities that also provide opportunities to interact, share knowledge, develop a common understanding, and collaborate.

3. Educate according to the dedicated education curriculum on IAS for schools.

4. Promote volunteer work in management projects.

    - Consider the diverse skills and motivations of volunteers and consider the engagement of different types of volunteers according to the specific task at hand; this may range from short, one-time assignments (e.g. playing an online game resulting in data for scientific evaluation) to long-term engagements (e.g. monitoring of invasive species population development by retirees or students).
    - Routinely integrate a citizen science component into IAS management projects that
        - is open, accessible, coordinated, collaborative and ethical;
        - provides participants with useful information, activities, feedback and acknowledgement (which also promotes further engagement);

  - makes use of up-to-date and innovative technologies integrated in open and user-friendly platforms (e.g. iNaturalist).
- Continuously monitor and regularly assess the usefulness and effectiveness of the different approaches for volunteer engagement in practice (e.g. regarding the type of activities, ways of communication, or the focal IAS) to improve the future use and long-term suitability of volunteer work in IAS management.

---

**Current situation:**

- IAS and management terminology is still used inconsistently.
- Public awareness of the benefits of using native/low risk species instead of introduced/high risk species (e.g. ornamental plants) is growing but still too low.
- Citizen science is gaining importance (e.g. see Citizen Science projects on IAS on EASIN website), but its full potential is still not used (Price-Jones *et al.* 2022); in particular, citizen science projects are often quite isolated from each other, which results in redundancy and missed opportunities for synergy effects.
- Conflicting ethical views in the general public and also among some conservationists on whether or not established IAS should be managed; in particular, killing organisms is often seen as unethical, even if the species are invasive.

### Biosecurity, monitoring, management and restoration

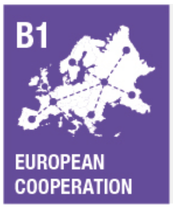


#### Goal B1:
Common effective biosecurity regime at the European and country level

*GOAL: By 2050, a common and effective biosecurity regime has been established in all European states (incl. non-EU states) and at the whole European level.*

**Actions to reach this goal:**

1. Apply and reinforce country-level horizon scans (HS) to guarantee high biosecurity within each country's borders.

   - Use standardized HS methodologies that can be used by the different countries but can still be adapted to the country's specific conditions.
   - Promote the exchange of information regarding HS/biosecurity between European countries, involving scientists in these dialogues, in order to: (i) avoid redundant efforts in different countries (e.g. consider IAS already identified as harmful in neighbouring countries), and (ii) enable capacity building in countries with low expertise.

2. Promote a European collaborative network between policy-makers, managers and researchers across countries to foster Europe-wide progress towards a common biosecurity regime (informed by HS).

   - Integrate such a network into available groups of experts (e.g. CBD, WHO, etc.).

- Use this network to encourage countries to follow recommendations and guidance from global networks/organizations (e.g. WTO, IMO) or other countries with high biosecurity expertise (e.g. Australia) for harmonizing biosecurity practices across Europe, in particular if binding tools of biosecurity coordination (e.g. intergovernmental agreements; see next action) are not possible to establish.
- Identify and explain the main advantages of a common biosecurity regime to stakeholders (standardized actions, easier implementation).

3. Prioritize, coordinate and monitor biosecurity measures through a European agency or intergovernmental agreement for IAS management and guided by an expert-based consultant group of managers, policy-makers and researchers from across Europe.

   - Identify and explain the main benefits of an intergovernmental agreement to stakeholders (e.g. efficient coordination of actions across countries, facilitation of monitoring and assessment).
   - Use the collaborative biosecurity network (see previous action) as a consultant organ.
   - Biosecurity priorities should be flexible and multi-layered to reflect different geographic opportunities: (i) continental level (common European priorities, e.g. species alien to entire Europe); (ii) biogeographical/regional level (priorities regarding species alien to part of Europe); (iii) country level (national priorities); and (iv) local level (local priorities).
   - If common biosecurity priorities cannot be established, at least establish common basic rules that all countries can agree on, especially regarding the overall goal of biosecurity efforts in each country (i.e. different methods can achieve similar goals).
   - If a European agency or intergovernmental agreement for IAS management is not possible to establish, the network created in the previous action should increase efforts to encourage exporting nations to implement biosecurity prior to export, e.g. through certification schemes, trade agreements or individual commodity negotiations.

---

**Current situation:**

- The IAS EU regulation (1143/2014) promotes that member states should make every effort to ensure close coordination among countries, but there is still a lack of communication and collaboration among Member States and between EU and non-EU countries.
- The free market within the EU (i.e. free movement of goods) counteracts effective invasion management and the enforcement of regulations.
- European countries face different IAS situations, so any biosecurity regime should consider these differences while still maintaining the overall perspective.

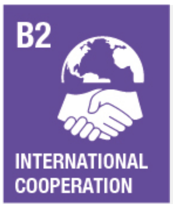


#### Goal B2:
Coordination of biosecurity regimes
between Europe and the rest of the world

*GOAL: By 2050, European biosecurity regimes are highly coordinated with those from other parts of the world that have strong connections (by trade and traffic) to Europe.*

**Actions to reach this goal:**

1. Establish a working group for coordination of biosecurity regimes at a global level, taking into account existing global networks/organizations (e.g. WTO, IMO). Tasks of the working group include:

    - Promote the coordination and harmonization of biosecurity regimes globally.
    - Facilitate the integration of relevant international organizations (e.g. related to food and human health) in biosecurity regimes, e.g. to certify or provide evidence of impact on biosecurity as well as to develop ways to incorporate biosecurity into trade agreements.

2. Learn lessons from countries with larger experience managing IAS (e.g. Australia, New Zealand, South Africa), identifying and analyzing (incl. cost assessment) novel management techniques that have proven to be useful and considering their applicability in the European context. Novel risks may stimulate novel responses and practices.

3. Strengthen the effective regulation of internal and external European borders, including the regulation of trade activities that represent introduction pathways to Europe and the alignment of biosecurity practices between European and non-European states.

    - Follow recommendations of global networks/organizations (e.g. WTO, IMO).
    - Consider mechanisms to control the movement of specific goods or species within the EU and Europe, including the opportunities provided by taking advantage of natural geographical barriers.
    - Gain support for stricter regulations by highlighting cases of economically important pests.

---

**Current situation:**

- Lack of or inefficient collaboration/communication between European (both EU and non-EU countries) and non-European countries.
- Biosecurity and conservation have different priorities on the political agenda of countries in different regions of the world.

B3

MONITORING AND PRIORITY SYSTEM

#### Goal B3:

Monitoring and assessing IAS

*GOAL: By 2050, a comprehensive regime for monitoring and assessing the introduction, establishment, spread and impacts of IAS has been established at the European level (incl. non-EU members) and the country level.*

**Actions to reach this goal:**

1. Foster collaboration among national agencies and stakeholders across Europe that have early-detection and assessment systems in place (i.e. integrated systems of active or passive surveillance to detect new IAS and assess their risk of establishment, spread and impact as early as possible so that decision-making can be done).
2. Identify pathways of introduction and spread of IAS, as well as areas occupied by IAS across European countries, following standardized protocols.
    - Identify the most promising and efficient ways to collate and homogenize available IAS data, and then develop or adopt relevant data standards (e.g. sources of information to use, how to store the data, formats, information to be exchanged between countries, a centralized data platform. Ideally, use a nested approach (e.g. Figure 3 in Latombe *et al.* (2017) that facilitates the integration of data across spatial scales.
3. Establish an early-detection and assessment system at the European and country level, especially in priority areas for monitoring.
    - Monitoring priorities can differ depending on the objective: areas where new introductions are likely (i.e. areas linked to pathways of introduction, e.g. ports or urban areas), or sensitive ecosystems of high biodiversity value where impacts are likely (e.g. protected area networks).
4. Promote the use of IAS apps for citizen science (targeted to different sectors or the general public) linked to early-detection and assessment systems, supported by up-to-date communication campaigns to show the benefits and importance of early-detection and assessment (Howard *et al.* 2022).
5. Secure long-term funding for the maintenance of early-detection and assessment systems.
    - Explain the importance of continuous surveys to companies and governments, e.g. reducing impacts, costs and time effort, ideally based on a cost-benefit analysis.

---

**Current situation:**

- Long-term monitoring of IAS and their impacts is not funded in Europe.
- We still cannot easily differentiate between pathways of primary introduction (e.g. intercontinental introductions to major ports) and of subsequent secondary introduction (e.g. intracontinental transport to smaller towns or natural spread of introduced species) to use limited management resources more efficiently.

#### Goal B4:
Sustained effort on minimizing entries of new IAS (prevention)

*GOAL: By 2050, sustained effort on minimizing the introduction of new IAS is secured (e.g. in terms of funding and personnel) in Europe for priority pathways.*

**Actions to reach this goal:**

1. Foster collaboration among national agencies and stakeholders across Europe that have rapid response systems in place (i.e., systematic effort to rapidly eradicate, contain, or control invasive species while the infestation is still localized).

    - Develop mechanisms to share and build on experience of rapid IAS eradication and control to develop long-term expertise, reduce costs and increase effectiveness (e.g. expert teams from multiple countries, dialogues among agencies, researchers and stakeholders etc.).
    - Coordinate the rapid eradication and control of IAS at the country level.

2. Build on or initiate exhaustive mapping of IAS distributions in each country to improve prevention, using standard protocols and recently updated data. If such data is not available, develop and apply workflows to standardize and integrate existing databases (e.g. Seebens et al. 2020) and use this information as a starting point to set preliminary priorities.

3. Identify and establish management priorities for IAS prevention, not only for species but also for areas and pathways (following McGeoch *et al.* (2016), see Figure 1 therein), at the European and country level.

    - To improve cost-effectiveness, assess the feasibility, costs and economic impacts of management while prioritising species, pathways and areas.
    - Consider synergies between existing management actions driven by economic requirements (e.g. keeping waterways clear) and the management of other IAS in the same systems.

4. Establish a rapid response system and infrastructure to reduce new introductions or isolated, new infestations of a previously established, invasive organism.

    - Use contingency planning and simulation exercises to test rapid response systems.
    - Provide data from management actions on a systematic and transparent/open data format to ensure maintenance and accessibility of knowledge.
    - Build on existing repositories (e.g. CABI Invasive Species Compendium) to create an open source platform that centralizes all information available on IAS management (LIFE projects, Nature 2000 activities, research papers, etc.).

5. Coordinate management responses for all priority pathways, to reduce the rate of new species arrivals.

6. Monitor and assess the achievement of pathway management objectives (at the European and country level), to improve management based on the monitoring/assessment outcomes using innovative approaches.

---

**Current situation:**

- EU IAS Regulation (1143/2014) requires prioritization and effective management of unintentional introduction pathways by all Member States.
- Few collaborations among European countries and between European and non-European countries in regard to pathway prioritization and management.

B5 CONTROL AND ERADICATION

#### Goal B5:

Sustained effort on IAS eradication and control

*GOAL: By 2050, sustained effort on eradicating and controlling IAS is secured (e.g. in terms of funding and personnel) in Europe*

**Actions to reach this goal:**

1. Foster collaborations between neighbouring countries facing similar IAS realities (e.g. Scandinavia countries, Germany & Austria) to harmonize management objectives across country borders so IAS populations can be managed consistently across jurisdictions;

2. Identify and establish management priorities for IAS eradication and/or control, not only for species but also for areas and pathways, at the European and country level.

    - Include an assessment of the feasibility and cost-effectiveness of eradication and control to inform prioritization for management, as well as a risk management analysis that assesses the risks connected to the management actions and the feasibility of reducing those risks (see Booy *et al.* 2017, 2020).
    - Recognise when IAS can no longer be cost-effectively managed at a population level and impact mitigation should be considered instead.
    - Link government subsidies (e.g. to farmers for the agri-environment) to receivers' commitment of undertaking effective IAS management and biosecurity.

3. Sustained effort in management of identified (established) IAS in priority areas.

    - Coordinate ongoing management actions on the ground, including activities that are not targeted to eradication or control of IAS but could if required (e.g. public gardening or forestry works managed by municipalities.
    - Promote volunteer work, both as people that dedicate time to it or people that do it in their properties.
    - Inform the public with clear instructions on how to proceed while carrying out management actions.

4. Sustained effort in management of identified (established) IAS beyond priority areas.

    - Highlight cross-sectoral interlinkages and emphasize the need to protect biodiversity to secure food provision and the good quality of life.

5. Monitor and assess the achievement of management objectives with respect to eradication and control (at the European and country level), to improve management based on the monitoring/assessment outcomes using innovative approaches.

    - Make use of tools to systematically compile evidence on the effectiveness of different management techniques (similar to e.g. the Conservation Evidence initiative), ideally in a centralized open data platform. Such tools should allow filtering the evidence by species/habitat/tool/geographic region to allow the user to consult the most relevant evidence available.

---

**Current situation:**

- Too little effort being devoted to practical management to achieve long-term goals.
- The management of widespread IAS receives much attention, but outcomes of such programmes for minimising the impact of IAS on biodiversity are often underreported.
- Management decisions are influenced by priorities (e.g. assets to be protected and further impacts to be avoided) and limited funds.
- There exists a conflict between investing in the removal of multiple species when prevention or complete removal is still feasible, or investing in expensive and often ineffective long-term management of a few species.

B6

RESTORATION

#### Goal B6:

Restoration activities

*GOAL: By 2050, restoration activities are undertaken in a way that is systematic and informed by research.*

**Actions to reach this goal:**

1. Foster communities of practice of researchers, managers and restoration practitioners and promote transdisciplinary collaboration between researchers in the field of invasion biology and restoration ecology (Wenger *et al.* 2002; Hulme 2006).

2. Undertake restoration activities in a systematic and sustained way, informed by research.

    - This includes activities to achieve biotic resistance and increased resilience of ecosystems, so that the impacts and spread of IAS are reduced, also tackling other anthropogenic disturbances (e.g. reduction of nutrient loads in a water body could help suppressing population of aquatic IAS plants without excessive long-term management costs).

- Regulations should incorporate 'polluter-pays' mechanisms that charge compensatory payment from stakeholders who undertake damaging or negligent activities in an area, and dedicate these payments for funding of restoration activities.
- Promote good practices of restoration (independently of the cause of degradation, be it IAS or other causes) that discourage the use or spread of IAS.
- Perform cost-benefit analyses of restoration activities, considering ecosystem services.

3. Monitor and assess the achievement of restoration objectives (at the European and country level), to improve restoration based on monitoring/assessment outcomes using innovative approaches.

   - Identify standard indicators to track recovery.
   - Promote public engagement to take advantage of widespread citizen science/volunteer monitoring.

---

**Current situation:**

- According to EU IAS Regulation (1143/2014), member States should carry out restoration measures to assist the recovery of ecosystems damaged by IAS.
- Restoration activities are not implemented systematically to recover degraded ecosystems by the impact of IAS.
- There is a lack of guidance, research on effectiveness and knowledge on how to restore ecosystems impacted by IAS.
